# Coral Probiotics Buffer Adjacent Ecosystem-Level Responses to Extreme Marine Heatwave

**DOI:** 10.64898/2026.03.30.715272

**Authors:** Yusuf C. El-Khaled, Francisca C. Garcia, Erika P. Santoro, Neus Garcias-Bonet, Matteo Monti, Marcos A.L. Teixeira, Micaela S.S. Justo, Gloria Gil-Ramos, Juan Sempere-Valverde, Glafira Kolbasova, Laura Beenham, Gustavo Duarte, Duarte Martins, Chakkiath Paul Antony, Torsten Thomas, Susana Carvalho, Raquel S. Peixoto

## Abstract

Probiotics can enhance coral thermal tolerance, yet their ecosystem-level effects remain unknown. Here, we present the first long-term *in-situ* test of whether coral-targeted probiotics influence adjacent cryptobenthic reef communities during a record marine heatwave. Probiotics were applied to *Pocillopora favosa* and *Acropora* spp. coral colonies for 18 months, spanning the fourth global bleaching event. Cryptobenthic communities were assessed using biomimetic monitoring structures integrating biodiversity surveys, molecular profiling, microbial network analyses, and metabolic assays. Before the heatwave, probiotic and control patches were comparable across structural, microbial, and functional metrics. Following thermal stress, control patches exhibited pronounced losses of cryptobenthic invertebrate abundance and taxonomic breadth, microbial network fragmentation, and net carbonate dissolution. In contrast, probiotic-treated patches retained higher biodiversity, cohesive microbial interaction architectures, and positive calcification. These findings demonstrate that coral-targeted probiotics can scale from host-level intervention to buffer adjacent ecosystem-level responses to extreme marine heatwaves under accelerating climate change.

**Teaser:** A coral-targeted probiotic strategy enhances multi-trophic resilience under heat stress.

## Main

Marine heatwaves are reshaping shallow marine ecosystems, triggering mass bleaching, habitat degradation, mass mortality of marine organisms - including but not limited to corals^>1– >3^, and restructuring of biogeochemical and food-web processes^1,4–6^. Their increasing frequency and duration indicate a transition from rare disturbances to recurrent environmental regimes^7,8^, exemplified by the fourth global coral bleaching event^9,10^.

Microbiome-based interventions such as Beneficial Microorganisms for Corals (BMCs) have emerged as promising tools to enhance resilience under extreme thermal stress^11–13^. Recent *in-situ* treatments indicate that probiotic applications can minimize the impacts of thermal stress and disease in corals^14,15^. Their broader influence on surrounding reef microbiomes and organisms is, however, only beginning to be evaluated^16–18^ and, thus, long-term, broader field assessments during natural marine heatwaves and evaluations of ecosystem-level consequences are lacking.

Cryptobenthic communities, commonly defined as diverse assemblages of sessile and motile invertebrates and associated microbes inhabiting reef cavities, contribute disproportionately to reef biodiversity, nutrient cycling, and productivity relative to their total biomass ^19,20^. Despite their functional importance, cryptobiomes remain rarely incorporated into reef monitoring frameworks. Given their short life cycles and sensitivity to environmental change, they may serve as early bioindicators of reef biodiversity and functional changes and broader disturbance impacts. Limited evidence indicates that bleaching and structural degradation of coral reefs can impact cryptobenthic assemblages ^21,22^, yet no studies have assessed how climate adaptation strategies affect cryptobiomes during real-world marine heatwaves. Here, we employed customized biomimetic Autonomous Reef Monitoring Structures (ARMS)^23^ to quantify cryptobenthic diversity, microbial organization, and ecosystem function, enabling assessment of ecosystem-level responses to microbiome-based interventions for corals during a natural marine heatwave.

Whether probiotic interventions applied to corals can directly or indirectly benefit surrounding reef habitats remains unknown. Cryptobenthic communities can contribute to post-disturbance recovery through multiple pathways, including rapid biomass turnover, nutrient regeneration that fuels primary production, and carbonate accretion by crustose coralline algae and other calcifiers that stabilize substrate and facilitate coral recruitment^24^. Disruption of these communities following marine heatwaves, subsequent coral bleaching and structural degradation can therefore impair both biogeochemical cycling and the physical processes underpinning reef recovery. Leveraging an 18-months *in-situ* coral probiotic treatment that encompassed a record marine heatwave in the central Red Sea (Fig. 1), we integrated biodiversity surveys, metabolic assays and microbial profiling and network analyses to test the extent of which effects can be quantified beyond the treated corals and persist through extreme thermal stress. By resolving cross-compartment responses, i.e., invertebrate and microbial diversity and stability, and cryptobiome functioning, our study addresses whether targeted microbiome interventions can operate at ecosystem-relevant scales, a key consideration for emerging climate adaptation strategies.

**Figure 1:**
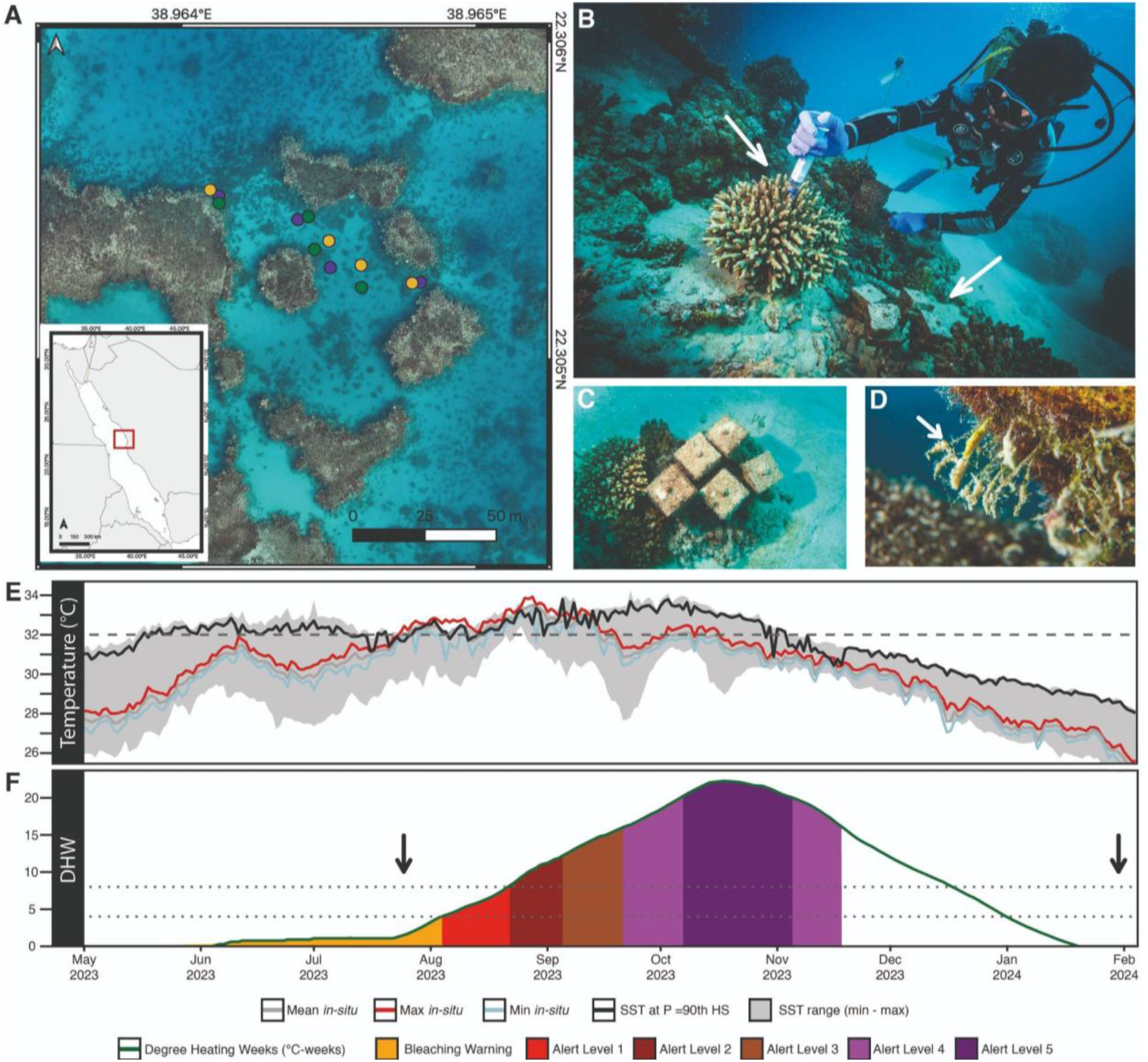
Study site, experimental deployment and thermal context. **(A)** Maps showing the location of the Coral Probiotic Village (CPV) and reef patches (Control: gold, BMC1: teal, BMC2: purple) within the study site located in the central Red Sea (red rectangle). **(B - C)** *In-situ* deployment and probiotic inoculation (white arrows) of biomimetic Autonomous Reef Monitoring Structures (ARMS). **(D)** Associated cryptobenthic invertebrate communities on ARMS (white arrow). **(E)** *In-situ* seawater temperatures (daily mean, maximum and minimum) recorded by a CTD logger, shown alongside satellite-derived sea surface temperatures (SST) including minimum, maximum and the NOAA 90th-percentile HotSpot SST (SST@90th HS). The dashed line at 32 °C marks a commonly used bleaching threshold. **(F)** Accumulated thermal stress shown as Degree Heating Weeks (DHW) with operational NOAA bleaching risk categories (Bleaching Alert Area, 7-day maximum) indicated as colored shading. The dotted lines mark 4 and 8 DHW, thresholds associated with bleaching risk, and arrows denote pre- and post-marine heatwave sampling time points. Photographs by Uli Kunz **(B–D)**.

### The 2023 marine heatwave as an ecosystem-wide disturbance

In 2023, coral reefs globally experienced extreme marine heatwaves culminating in the fourth global bleaching event^9,10^. Within this global context, the central Red Sea faced a record marine heatwave, with *in-situ* daily mean seawater temperatures at the study site reaching 33.05 °C at 6.5 - 8 m depth (Fig. 1E). Cumulative thermal stress peaked in October 2023, with relative Degree Heating Weeks (rDHW) reaching 17.57 °C-weeks (daily mean) and 21.79 °C-weeks (daily maximum)^25^, consistent with NOAA-reported regional maxima of 22.26 DHW (Fig. 1F). This event coincided with widespread coral bleaching across the region and served as a natural disturbance separating pre- and post-heatwave sampling periods.

### Probiotic treatments buffer cryptobenthic community structure during extreme thermal stress

To assess whether coral-targeted probiotics influence adjacent reef communities, we established replicated coral-centered reef patches consisting of individual treated coral colonies and adjacent customized biomimetic Autonomous Reef Monitoring Structures (ARMS) positioned 10-60 cm away from inoculated corals. Two native Beneficial Microorganisms for Corals (BMC) consortia (BMC1 as described in ref.^16^, and BMC2 as described in ref.^26^) were applied to reef-building corals three times per week over an 18-month period encompassing the marine heatwave. Briefly, probiotic consortia were resuspended in a saline solution (3.5% NaCl) and were applied directly onto coral colonies (BMC1 on *Pocillopora favosa*, BMC2 on *Acropora* spp.) using a syringe as described in ref.^16^, whereas ARMS were not inoculated, allowing assessment of indirect ecosystem-level effects on adjacent cryptobenthic assemblages (Fig. 1B-D). More details about the application of probiotics and the experimental design are available in the online methods and supplementary material.

Prior to the heatwave, visually identified motile invertebrate assemblages were dominated by Annelida, Arthropoda, and Mollusca, with comparable richness and abundance across control and probiotic-treated patches (generalized linear models, all *p* > 0.9; Fig. 2A, 3A; Supplementary Table S1), indicating no detectable off-target effects of probiotic application under ambient conditions. However, following the marine heatwave, visually identified cryptobenthic macrofaunal abundance declined across all treatments, with the decline being markedly more severe in control (placebo) patches (Fig. 3A; Supplementary Tables S1 and S2). Control assemblages experienced a significant post-heatwave reduction in both mean abundance (-82.9 ± 8.6 %; post hoc contrasts on estimated marginal means *p* = 0.009; Fig. 3A; Supplementary Table S1 and S2) and phylum-level richness (Fig. 2), while BMC2 treated patches retained significantly higher phylum-level richness relative to controls (generalized linear models with Tukey-adjusted post hoc contrasts, *p*_adj_ = 0.031) and BMC1 followed same direction but showing a weaker effect (*p*_adj_ = 0.069); Supplementary Table S1). Post-heatwave, both probiotic treated cryptobenthic communities displayed approximately three- to four-fold higher abundances of motile cryptobenthic invertebrates compared to controls (post hoc contrasts on estimated marginal means, *p* < 0.05, Supplementary Table S2), consistent with a treatment-associated buffering effect. Notably, these treatment- and time-dependent differences were not accompanied by changes in body size structure: community-weighted mean size of motile cryptobenthic invertebrates did not change significantly between sampling points within any treatment (two-way linear model with planned EMM contrasts, all *p* > 0.36; Supplementary Table S3), indicating that observed responses reflected shifts in abundance and structure rather than size-selective effects.

**Figure 2:**
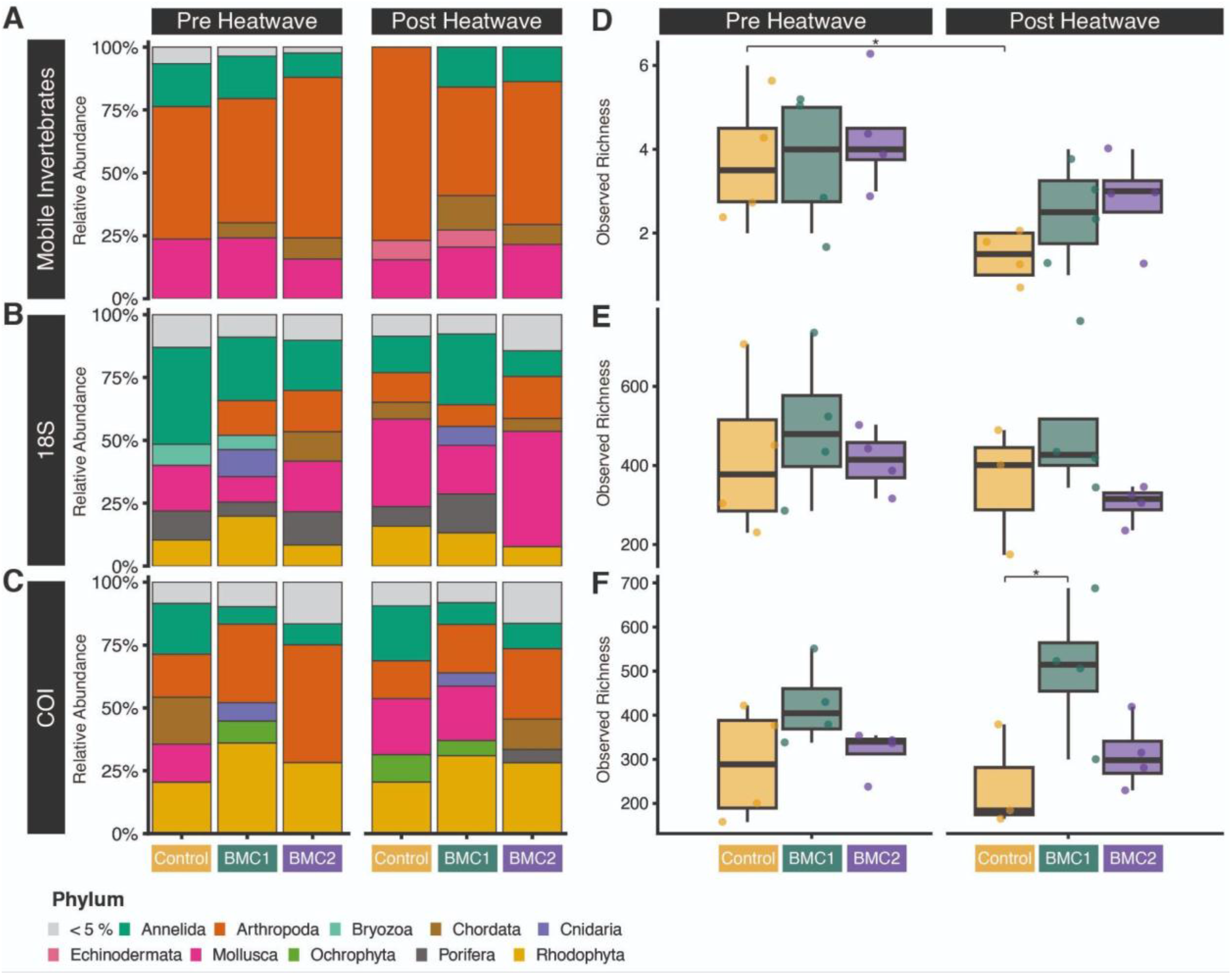
Shifts in community composition and richness following the marine heatwave. Stacked bar plots show relative phylum-level composition of reef patches receiving Control (placebo), BMC1, or BMC2 treatments before (pre) and after (post) the 2023 marine heatwave (MHW). Panels represent **(A)** visually identified mobile invertebrates, **(B)** 18S metabarcoding, and **(C)** COI metabarcoding. Bars are grouped by sampling period and treatment, with colors indicating phylum-level relative abundance. Control patches exhibited pronounced post-heatwave compositional shifts and reduced phylum representation of mobile invertebrates, whereas probiotic-treated patches maintained comparatively broader taxonomic representation, particularly in the 18S dataset (see Supplementary Table S4 for PERMANOVA results). Patterns were directionally consistent between molecular and visual datasets and align with higher cryptobenthic abundance and crustose coralline algae (CCA) cover observed in probiotic-treated patches (Fig. 3; Supplementary Tables S1-S6). **(D-F)** Corresponding observed richness reflects the number of taxa (visual surveys) and ASVs (18S and COI marker genes) per patch. Asterisks denote statistically significant planned contrasts based on estimated marginal means (Tukey-adjusted for treatment comparisons within time, *p* < 0.05; see Supplementary Table S6 for full results). Post-heatwave communities exhibited compositional reorganization and treatment-dependent shifts in richness.

**Figure 3:**
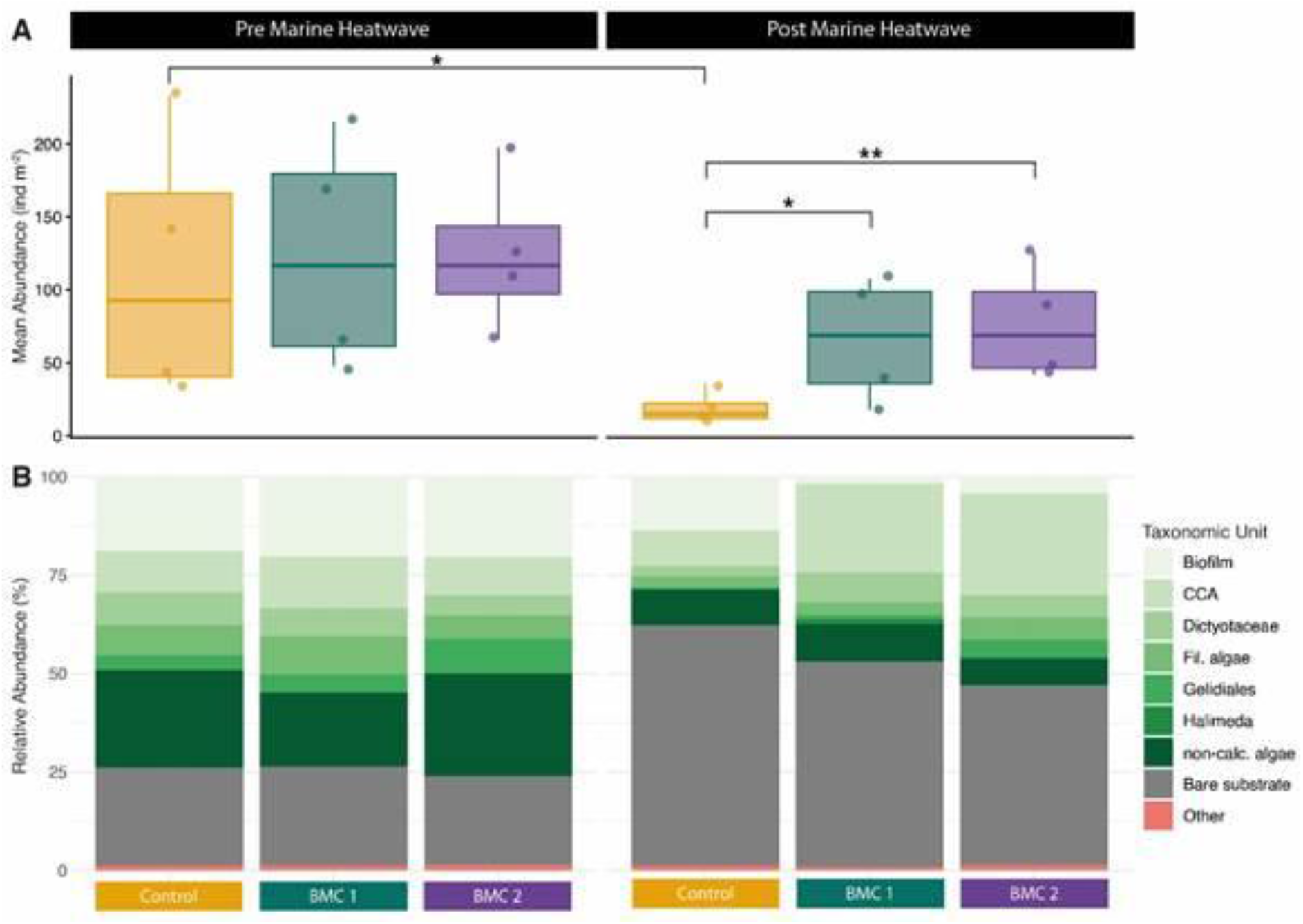
Probiotic treatments mitigate heatwave-driven collapse of visually identified cryptobenthic assemblages. **(A)** Abundance of motile cryptobenthic invertebrates (mean ± SE) before and after the heatwave. Abundances declined across all treatments, but declines were 3-4-fold greater in controls (pre-vs post-heatwave ratio = 5.846, Tukey-adjusted p = 0.009; Supplementary Table S1), corresponding to a 82.9 ± 8.6% reduction (Supplementary Table S2). Post-heatwave, probiotic treated patches retained 3-4-fold higher abundances than controls (Control vs BMC1 ratio = 3.385 ± 1.281, *p* = 0.046; Control vs BMC2 ratio = 3.923 ± 1.319, *p* = 0.008; Supplementary Table S1). Body-size distributions did not change across treatments or years (*p* > 0.36; Supplementary Table S3), indicating that responses reflected abundance loss rather than size-selective filtering. **(B)** Sessile ARMS assemblages showed increased bare substrate and microbial biofilm cover in controls, whereas probiotic-treated patches retained higher crustose coralline algae (CCA) cover (pairwise Wilcoxon tests, BH-adjusted *p* ≤ 0.1; Supplementary Table S7). Together, these patterns indicate treatment-associated buffering of both motile and sessile cryptobenthic structures under extreme thermal stress.

Molecular surveys broadly corroborated visual patterns. Sequential PERMANOVA (Type I sums of squares) on Bray-Curtis dissimilarities revealed significant effects of sampling time for both eukaryotic assemblages characterized by 18S rRNA gene amplicons (R^2^ = 0.068, *p* = 0.027) and metazoan assemblages derived by COI amplicons (R^2^ = 0.066, *p* = 0.004), indicating heatwave-associated shifts in multivariate community composition (Supplementary Table S4). Treatment explained additional variation in 18S-derived eukaryotic assemblages (R^2^ = 0.119, *p* = 0.027) and showed a marginal effect in COI-derived metazoan assemblages (R^2^ = 0.101, *p* = 0.079), whereas time × treatment interactions were not significant for either marker (18S: *p* = 0.224; COI: *p* = 0.331). Tests of homogeneity of multivariate dispersion were non-significant for all comparisons (PERMDISP, all *p* > 0.32; Supplementary Table S4), indicating that PERMANOVA results were not driven by differences in within-group dispersion. Alpha diversity metrics derived from both molecular markers (observed richness, Shannon diversity, Simpson diversity, Chao1 richness) showed no significant time × treatment interactions in two-way linear models (Type II ANOVA; Supplementary Table S5). However, COI-derived metazoan assemblages exhibited significant main effects of ‘treatment’ independent of ‘time’ (Observed: F = 6.495, *p* = 0.008; Chao1: F = 6.721, *p* = 0.007; Supplementary Table S5). Planned post hoc contrasts using estimated marginal means (Tukey-adjusted within ‘time’; Supplementary Table S6, Supplementary Fig. S1) revealed lower post-heatwave richness in control relative to BMC1 patches (Observed: *p*_adj_ = 0.017; Chao1: *p*_adj_ = 0.016). Together, these results indicate that heatwave-driven diversity loss was strongest in visually surveyed assemblages, whereas treatment effects in metabarcoding datasets were expressed as among-treatment differences rather than divergent temporal trajectories, which is consistent with the greater post-heatwave decline in diversity observed in control patches.

Comparable patterns were observed in the photoautotrophic cryptobenthic fraction. Prior to the heatwave, no detectable differences on the photoautotrophic cover and substrate composition were observed between treatments (Supplementary Table S7 and S8). Following the heatwave, univariate analyses indicated shifts in specific functional categories rather than whole-community restructuring. In post-heatwave assemblages, control ARMS exhibited higher relative cover of microbial biofilms and lower crustose coralline algae (CCA) cover compared to probiotic-treated patches (Kruskal-Wallis raw *p* ≤ 0.05; BH-adjusted pairwise Wilcoxon *p* = 0.078; Supplementary Table S7). However, multivariate analysis of photoautotrophic and sessile assemblage composition using Bray-Curtis dissimilarities did not detect a significant overall treatment effect (PERMANOVA R^2^ = 0.19, *p* = 0.516), and dispersion did not differ among treatments (PERMDISP *p* = 0.590, Supplementary Table S8). These results indicate that probiotic-associated differences were expressed primarily through changes in specific photoautotrophic categories, particularly retention of CCA, rather than broad shifts in sessile community composition.

### Probiotic treatments preserve cryptobenthic microbial community networks during heat stress

Bacterial communities (characterized using 16S rRNA gene amplicon sequencing) exhibited modest heatwave-associated changes but no treatment specific-turnover. Alpha diversity was quantified at the ASV level, i.e., observed ASV richness, Shannon and Simpson diversity, and Chao1 richness, and declined slightly across time independent of treatment (Supplementary Table S5), and compositional analyses detected no significant time × treatment interactions, indicating an overall heatwave-associated reduction in within-sample bacterial diversity associated with thermal stress. Multivariate analyses based on Bray-Curtis dissimilarities detected a significant overall model effect (PERMANOVA R^2^ = 0.325, *p* = 0.004, Supplementary Table S9A). However, pairwise contrasts revealed no significant differences among treatments within either sampling point (pre-heatwave *p* = 0.356; post-heatwave *p* = 0.919), and temporal shifts were significantly only within BMC2-treated patches (*p* = 0.025; Supplementary Table S9B). Importantly, when community composition was analyzed using Aitchison distances (CLR-transformed data), no significant time × treatment interaction was detected (*p* = 0.30), indicating that treatment-specific restructuring was not supported when accounting for the compositional nature of amplicon data. Homogeneity of multivariate dispersion was confirmed for all models.

Aligning with these results, differential abundance testing using ANCOM-BC2 identified no ASVs that were significantly differentially abundant among treatments or across the heatwave after false discovery rate correction (FDR ≤ 0.05; Supplementary Table S10). Across all contrasts, hundreds to thousands of ASVs were tested, yet none exhibited statistically robust shifts in abundance. Family-level relative abundance profiles similarly showed no systematic restructuring across treatments or time points (Supplementary Fig. S2). Moreover, the bacterial taxa comprising the applied BMC consortia were not detected in ARMS-associated 16S datasets at either time point, indicating that observed community and network responses were not driven by direct incorporation of probiotic strains into cryptobenthic assemblages. Together, these results indicate that bacterial responses to the marine heatwave were modest at the taxon-abundance level and not detectably influenced by probiotic treatment. No statistically supported treatment-specific taxonomic turnover was detected. As such, probiotic-associated effects on cryptobenthic bacterial communities were not expressed through shifts in ASV-level composition, motivating subsequent analyses of higher-order interaction structure.

To assess microbial interaction restructuring independent of taxonomic turnover, we inferred co-occurrence networks using the 200 most abundant ASVs present in at least two samples. This standardized feature set captured >83% of total sequencing reads across treatments and sampling periods and ensured comparability across networks (Supplementary Fig. S3). This approach ensured that all networks were constructed from a shared feature space and minimized artefacts associated with condition-specific rank shifts or sparsely observed taxa, while still retaining the dominant fraction of the microbial signal.

Prior to the marine heatwave, microbial networks were similarly organized across all treatments. Each microbial network comprised a single, dominant connected component with comparable edge density and modularity (edge density 0.34-0.37, modularity 0.35-0.41; Fig. 4A, C and E), indicating broadly conserved interaction architectures prior to the stress event.

**Figure 4:**
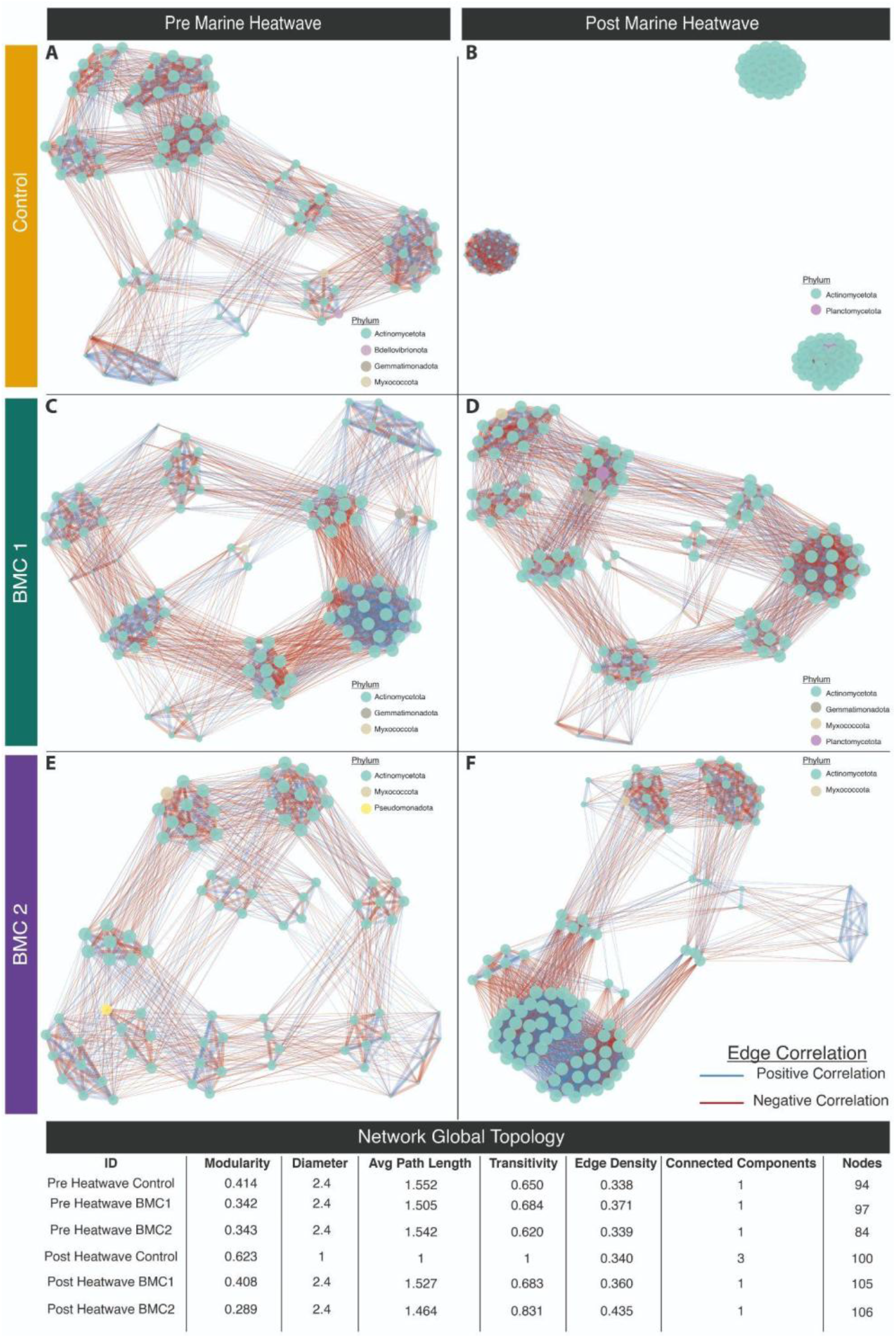
Probiotic treatments modulate microbial interaction-network responses during an extreme marine heatwave. Meta-community co-occurrence networks (top 200 ASVs present in ≥2 samples, representing >83% of sequencing reads) are shown for control **(A-B)**, BMC1 **(C-D)**, and BMC2 **(E-F)** patches before **(A, C, E)** and after **(B, D, F)** the heatwave. Control networks exhibited pronounced restructuring following the heatwave, including reduced edge overlap (Jaccard similarity *J* = 0.098), increased compartmentalization (one to three connected components), and shifts in global network metrics relative to bootstrap expectations, consistent with loss of cohesion. Networks from BMC1-treated patches retained comparable global topology across time, with no significant pre-post heatwave differences in edge density or modularity based on permutation tests. Networks from BMC2-treated patches showed directional changes in global metrics (increased edge density, decreased modularity), but permutation-based contrasts did not support statistically significant differences in overall topology. Edge-level analyses indicated widespread changes in association strength in probiotic-treated networks, whereas no individual associations changed significantly in controls after false discovery rate correction. Centrality distributions shifted significantly in control and probiotic networks, indicating redistribution of node roles under heat stress. See Supplementary Table S11 and online repository for full network metrics, permutation tests, and bootstrap analyses.

Following the marine heatwave, network trajectories diverged strongly among treatments. Networks associated with probiotic-treated patches largely retained cohesive structures. In BMC1 patches, global network topology remained effectively unchanged between pre- and post-marine heatwave conditions, with no significant differences in edge density or modularity detected by permutation-based network comparisons (all permutation tests *p* ≥ 0.25; Fig. 4C and D; Supplementary Table S11). BMC2 networks exhibited a marked though non-significant shift in global topology, characterized by increased edge density and a concomitant decrease in modularity (Δedge density = 0.096 permutation *p* = 0.335; Δmodularity = -0.159; permutation *p* = 0.340), consistent with controlled reconfiguration toward a more integrated interaction structure rather than network fragmentation (Fig. 4F, Supplementary Table S11). Notably, while raw global metrics (e.g., edge density and modularity) can be compared descriptively, formal inference on topology differences was based on permutation contrasts (label-shuffling) reported in Supplementary Table S11. In contrast, microbial networks associated with control patches underwent pronounced structural disruption following the heatwave. Although constructed from an identical set of ASVs, pre- and post-heatwave control networks exhibited low edge overlap (Jaccard similarity *J* = 0.10), indicating extensive rewiring of microbial associations within a conserved node set (Fig. 4A and B; Supplementary Table S11). To test whether these changes could be explained by sampling variability alone, we generated a bootstrap null distribution by repeatedly reconstructing networks from resampled pre-heatwave control communities (k = 3, with replacement; 300 iterations). Across multiple independent metrics, including edge number, edge density, connectivity, and modularity, post-heatwave control networks occupied the extreme tails of the null distributions (empirical one-tailed *p* ≤ 0.001 for all metrics; Supplementary Table S11), indicating that the observed structural changes exceed expectations from stochastic resampling alone. Together, these patterns indicate that extreme thermal stress was associated with system-level disorganization of microbial interaction structure in control patches, whereas probiotic-treated communities maintained cohesive or directionally reconfigured network architectures.

At the level of individual associations, no individual pairwise association between ASVs exhibited a significant change in strength between pre- and post-heatwave control networks after false discovery rate correction, indicating that network reorganization arose from distributed, system-level shifts rather than isolated rewiring events. Node-level analyses supported these patterns: eigenvector centrality distributions shifted markedly in control networks across the heatwave (permutation *p* < 0.001), reflecting redistribution of node importance within a reorganized interaction network. Significant centrality shifts were also detected in BMC1 and BMC2 networks (both *p* < 0.001), but occurred in the absence of global network disruption in BMC1 and in the context of controlled reconfiguration in BMC2 (Supplementary Table S11).

Together, these results demonstrate that probiotic application was associated with distinct microbial interaction-network responses to extreme thermal stress. Whereas control communities underwent widespread system-level reorganization, BMC1 networks largely preserved pre-disturbance interaction architecture despite shifts in node centrality, and BMC2 networks exhibited controlled reconfiguration toward a more integrated topology without fragmentation. Importantly, these patterns persist under a conservative bootstrap framework that controls for sampling variability and network definition even using all ASVs present in at least two samples, indicating that the observed treatment effects reflect genuine differences in microbial interaction dynamics rather than artefacts of data sparsity or feature selection (Supplementary Fig. S4). Repeating the network inference using all ASVs present in at least two samples yielded qualitatively similar treatment-dependent trajectories, albeit with increased stochasticity and reduced resolution of global metrics, consistent with the expected influence of sparse, low-abundance features on correlation-based networks.

### Probiotic treatments preserve carbonate production despite reduced metabolism

The 2023 marine heatwave was associated with an overall tendency toward lower metabolic rates across sampling periods, with net and gross photosynthesis (P_net_ and P_gross_, respectively) generally reduced post-heatwave, while dark respiration (R_dark_) remained comparatively stable (Fig. 5; Supplementary Table S12). However, statistically significant temporal declines were detected only in BMC1-treated assemblages (P_net_: *p*_adj_ = 0.004, P_gross_: *p*_adj_ = 0.025), whereas control and BMC2 patches showed no significant pre-post changes. Importantly, no significant differences among treatments were detected within either sampling period for any metabolic flux (Supplementary Table S12). Together, these results indicate that primary productivity exhibited modest temporal variation but no detectable treatment effect.

**Figure 5:**
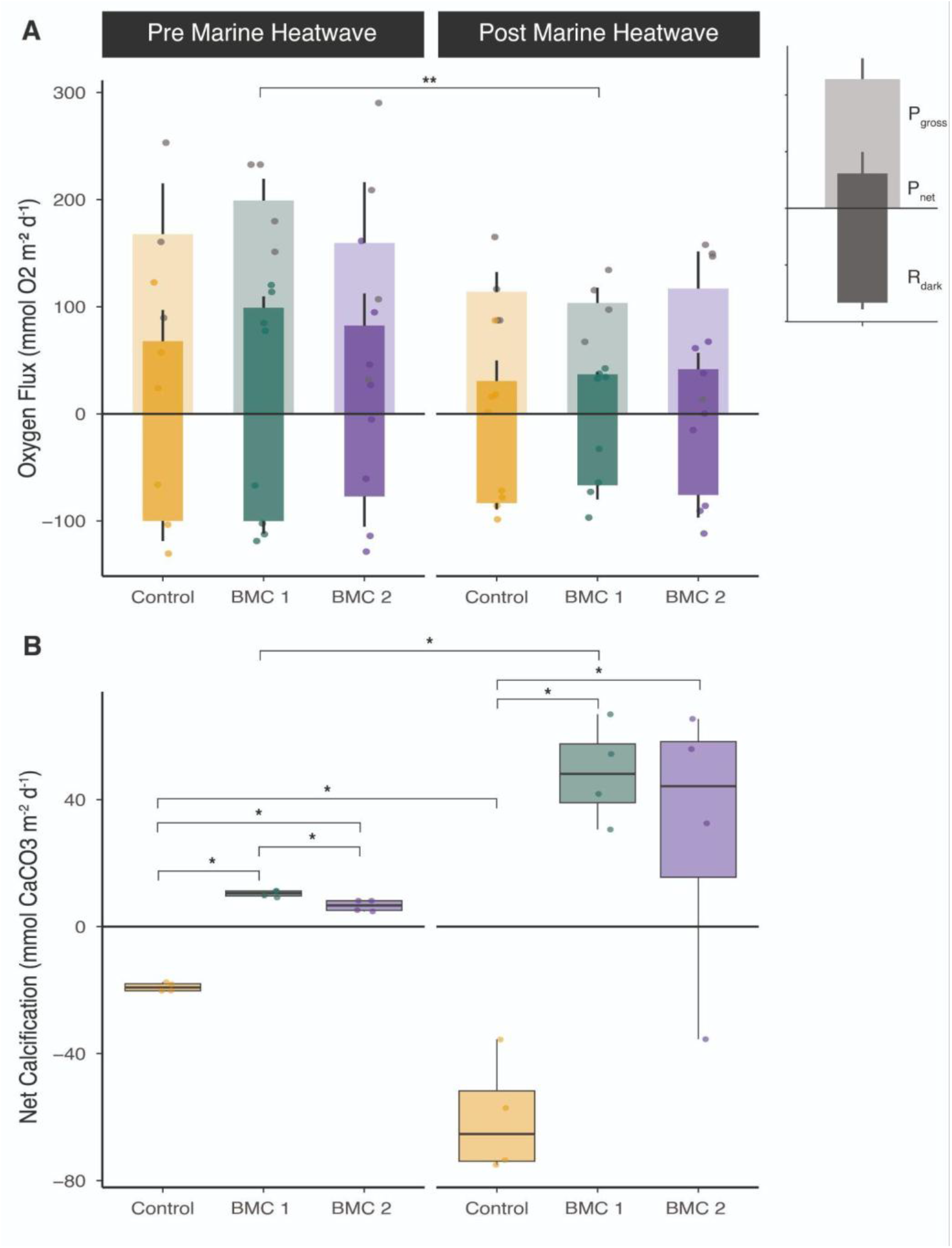
Probiotic treatments preserve carbonate accretion despite reduced metabolism. **(A)** Oxygen fluxes (gross photosynthesis, net photosynthesis, and dark respiration) measured prior to and after the marine heatwave. Significant temporal declines in net and gross photosynthesis were detected only in BMC1 treated patches (*p*_adj_ = 0004 and 0.025, respectively). No significant differences among treatments were detected within sampling periods. **(B)** Net calcification diverged sharply among treatments: controls exhibited net dissolution that intensified following the heatwave (*p*_adj_ = 0.043), whereas BMC1-treated patches showed a significant increase in calcification across time (*p*_adj_j = 0.043). BMC2 patches maintained positive calcification without significant temporal change. Controls had significantly lower calcification than both probiotic treatments before and after the heatwave (pre: *p*_adi_ = 0.029, post: *p*_adi_ = 0.043), whereas BMC1 and BMC2 did not differ post heatwave. Data shown as means ± SE overlaid with boxplots. Full statistical results in Supplementary Tables S12 and S13.

In contrast to metabolic rates, cryptobenthic carbonate dynamics diverged strongly among treatments. Control patches showed net dissolution both before and after the heatwave, with dissolution intensifying following the marine heatwave (t-test, *p*_adj_ = 0.043; Supplementary Table S13A). In contrast, probiotic-treated patches maintained positive calcification throughout the study period. Prior to the heatwave, calcification rates in both BMC treatments were significantly higher than in controls (ANOVA with Tukey’s HSD, *p*_adj_ = 0.029, Supplementary Table S13B), and this difference persisted post-heatwave (ANOVA with Tukey’s HSD, *p*_adj_ = 0.043). Following the heatwave, BMC1-treated patches showed a significant increase in calcification across the disturbance period (*p*_adj_ = 0.043), whereas BMC2-treated patches displayed no significant temporal change (*p*_adj_ = 0.343; Fig. 5; Supplementary Table S13). Across patches, net calcification rates were not significantly correlated with CCA cover or pooled calcifier abundance (all *p* > 0.3; Supplementary Table S14), indicating that carbonate fluxes were not explained by calcifier cover alone.

Together, these results demonstrate that although metabolic rates showed no treatment-specific effects, probiotic application to adjacent corals is associated with the preservation, and, in case of BMC1 enhanced, carbonate accretion, a key function underpinning reef framework stability and post-disturbance recovery.

## Discussion

Our findings demonstrate that coral-targeted probiotic interventions applied *in-situ* and tracked over 1.5 years can generate ecosystem-level effects that extend beyond treated hosts. After a record marine heatwave, untreated reef patches experienced sharp declines in cryptobenthic invertebrates, microbial network fragmentation, and net carbonate dissolution, whereas probiotic-treated patches retained higher biodiversity, cohesive microbial interaction architectures, and remained positive net calcification. These consistent changes across community structure, microbial interaction networks, and ecosystem function indicate that microbiome-based interventions can buffer and protect adjacent reef cryptobenthic communities after extreme thermal stress - a type of disturbance expected to become more frequent and severe under future warming scenarios^7,8,27^.

Importantly, these ecosystem-level differences emerged without detectable off-target effects prior to the heatwave. Under ambient conditions, probiotic- and control-treated patches exhibited comparable cryptobenthic composition, microbial diversity, network structure, and metabolic rates, consistent with natural levels of spatial heterogeneity previously described for cryptobiomes in the Indo-Pacific and Red Sea^22,28,29^. This indicates that treatment-dependent trajectories after the heatwave reflect differential responses to disturbance rather than detectable pre-existing differences among reef patches.

The 2023 marine heatwave acted as an ecological filter on cryptobenthic assemblages, but responses diverged sharply across treatments in both visual surveys and metabarcoding data. Control patches experienced substantial reductions in motile invertebrate abundance and phylum-level representation, alongside increased bare substrate and microbial biofilm cover. Such shifts are consistent with the sensitivity of cryptobenthic assemblages to structural degradation and microhabitat loss following coral bleaching^21,30^. In contrast, probiotic-treated patches retained broader taxonomic representation, higher abundances of motile fauna, and greater CCA cover. Given the role of CCA in substrate stabilization, carbonate accretion, and coral recruitment^31–33^, this retention likely supports post-disturbance recovery trajectories. Body-size distributions remained unchanged, indicating that buffering operated through preservation of taxonomic breadth and abundance rather than size-selective survival.

In contrast to macrofaunal and photoautotrophs responses, bacterial taxonomic structure based on 16S marker genes showed only modest heatwave-associated shifts and no detectable treatment-specific turnover. Alpha diversity declined slightly across time independent of treatment, and Aitchison-based structural analyses did not support significant treatment-by-time interactions. Differential abundance testing likewise detected no ASVs that were significantly enriched or depleted among treatments. However, despite this apparent taxonomic stability, microbial interaction structure diverged markedly. In control (placebo) patches, microbial co-occurrence networks became extensively rewired and fragmented following the heatwave, with reduced cohesion and increased modularity - patterns consistent with declining coordination and robustness under environmental stress^34,35^. In contrast, probiotic-treated patches retained their pre-heatwave topology, while BMC2 networks reorganized without collapsing into disconnected components. These findings indicate that probiotic-associated buffering was not expressed through shifts in microbiome composition, but through preservation or controlled restructuring of interaction networks. Network-level stability despite taxonomic constancy suggests that microbial resilience under disturbance may depend more on the organization of associations than on wholesale compositional turnover. Such system-level reorganization without differential abundance is increasingly recognized as a feature of microbial community responses to environmental change^36^.

Functional trajectories mirrored these structural and microbial patterns. Cryptobenthic metabolism declined across all treatments between sampling periods, likely reflecting a combination of heatwave disturbance and seasonal cooling. In contrast, results for carbonate accretion diverged strongly: control patches, already net decalcifiers prior to the heatwave, experienced further net dissolution, whereas probiotic-treated patches maintained positive calcification and, in the case of BMC1, increased carbonate production. Because CCA and other calcifiers contribute to carbonate accretion and substrate stability^31,37^, the higher retention of CCA in treated patches is consistent with preserved and enhanced net calcification. However, calcification rates were not linearly correlated with CCA or pooled calcifier abundance, indicating that carbonate fluxes likely reflect the integration of multiple processes, including dissolution, microenvironmental chemistry, and microbial mediation^38,39^, rather than CCA cover alone. The decoupling of metabolism and calcification suggests that functional resilience emerged from preserved community structure rather than sustained productivity alone.

Several non-mutually exclusive mechanisms may underlie these patterns, including altered microhabitat conditions around treated corals, stabilization of microbial interaction scaffolds, and reduced dysbiosis or pathogen propagation across adjacent substrates^36,38–41^. Notably, BMC strains were not detected on ARMS-associated communities, supporting an indirect mode of action in which coral-targeted probiotics modulate local microbial dynamics without establishing in off-target substrates. Importantly, probiotics do not need to colonize the host to trigger beneficial microbiome restructuring and holobiont function^42,43^. Although causal pathways remain unresolved, these cross-compartmental feedbacks illustrate how host-targeted microbiome interventions can influence ecosystem processes at adjacent micro-spatial scales.

Together, these findings extend reef resilience beyond coral hosts to the coupled dynamics of microbes, and their role supporting macroorganisms and underlying ecosystem functioning^44^. Rather than driving large taxonomic shifts, probiotic application was associated with preservation of interaction structure and carbonate accretion after extreme heat stress. This aligns with emerging “One Reef Health” perspectives emphasizing that resilience to climate extremes arises from integration across biological compartments rather than isolated components^39,45–48^, which is especially triggered by microbes^48^, and supports a “reef dysbiotic cascade” mechanism. Stabilizing the coral holobiont could therefore mitigate downstream ecological disruption across adjacent microhabitats, thereby indirectly buffering cryptobenthic assemblages.

Coral probiotics may act as an ecosystem-relevant tool capable of buffering climate-driven disturbance beyond individual hosts. While such interventions cannot substitute for rapid greenhouse gas emissions reductions, active restoration and rehabilitation will be required to maintain coral populations while carbon neutrality is pursued^49^. Within this broader adaptation portfolio, probiotics can complement such restoration and management strategies by enhancing local resistance to extreme warming events, allowing for adaptation and extended resilience and sustaining key functional currencies - including carbonate accretion - that underpin reef recovery.

## Figures

## Online Methods (short version, extended version in Supplementary Material - see below)

Extended versions of all methods used in this study, including detailed descriptions, parameters, and effect sizes and statistical approaches are provided in Supplementary Material.

### Study system and experimental design

The study was conducted in the *Coral Probiotic Village* in the central Red Sea (Northern Al Fahal reef complex, Saudi Arabia; 22.18°N, 38.96°E; Fig. 1A) between July 2022 and January 2024. Environmental parameters including seawater temperature, light, dissolved oxygen, and currents were continuously monitored using CTD, PME miniDOT, and ADCP sensors as described in Garcias-Bonet et al. (2025)^25^. Twelve discrete reef patches (2-3 m^2^, 5-8 m depth) were selected, each containing at least one *Pocillopora favosa* and *Acropora* spp. colony, sponges, natural reef matrix, and sandy substrate. Customized biomimetic limestone ARMS consisting of two horizontally stacked tiles on stainless steel rods were deployed 10-60 cm away from inoculated corals to investigate cryptobenthic communities (Fig. 1B).

### Probiotic treatments

Two native probiotic consortia (Beneficial Microorganisms for Corals; BMCs) and a control (placebo) consisting of a saline solution were applied to two coral species *in-situ* three times per week for 18 months. BMC1 (tailored to *P. favosa*) comprised six bacterial strains (*Halomonas piezotolerans, Bacillus aequororis*, two *Pseudoalteromonas* sp., and two *Cobetia* sp.), and BMC2 (tailored to *Acropora* spp.) comprised four strains (*Halomonas piezotolerans, Cobetia* sp., *Pseudoalteromonas* sp., *Pseudoalteromonas lipolytica*). These stains are part of the KAUST biobank and were previously isolated from Red Sea corals and screened for beneficial traits^16^. Per application, coral colonies received 30 mL of a 10^9^ cells mL^−1^ suspension, while the other coral species in the same patch received a saline solution as a control. In control patches, both species received a saline solution only mimicking the physical application using syringes. Treatments were assigned at the patch scale (n = 4 patches per treatment). Importantly, ARMS did not receive direct probiotic treatment (Fig. 1B).

### Sampling timeline and heatwave context

Biomimetic ARMS were retrieved after 12 months (July 2023; hereafter referred to as ‘pre-heatwave’ or variations thereof) and 18 months (January 2024; hereafter referred to as ‘post-heatwave’ or as variations thereof). *In-situ* seawater temperatures and NOAA Coral Reef Watch Virtual Stations confirmed that 2023 corresponded to a record heatwave for the region, peaking at >21 °C-weeks relative Degree Heating Weeks and accompanied by widespread coral bleaching. Pre- and post-heatwave incubations were performed at *in-situ* temperatures (∼30 °C and ∼25-26 °C, respectively). Postprocessing of the ARMS followed standardized approaches described in Leray and Knowlton (2015)^23^.

### Cryptobenthic community structure

Mobile invertebrates were visually identified, counted and size of each individual were measured following ARMS protocols^23^ (Leray and Knowlton 2015). Sessile and photoautotrophic taxa were quantified through photographic analysis with stratified point counts (40 per photo) to estimate autotroph cover and assemblage composition. Taxa were identified to the lowest feasible level, and sizes of mobile invertebrates were recorded. Phylum-level composition and diversity metrics were calculated for visual datasets.

### Metabolic and carbonate fluxes

Community metabolic fluxes were quantified via closed-cell incubations using a CISME device (Qubit Systems, Canada)^50^. Dark respiration and net and gross photosynthesis were calculated from continuous dissolved oxygen measurements. Discrete water samples were used to derive changes in total alkalinity (TA) and estimate net calcification/dissolution assuming a 2:1 TA:CaCO_3_ stoichiometry^51^. Fluxes were normalized to ARMS surface area and blank-corrected. All fluxes are displayed in mmol O_2_ or CaCO_3_ m^-2^ d^-1^, respectively.

### Molecular profiling

Genomic DNA was extracted from homogenized ARMS scrapings using PowerSoil-based protocols and subjected to amplicon sequencing targeting the bacterial 16S rRNA gene (V3-V4 region) and the eukaryotic 18S rRNA (V4 region) and COI marker genes. Libraries were prepared following standard Illumina protocols and sequenced on a NovaSeq 6000 platform (paired-end 250 bp). Amplicon sequence variants (ASVs) were inferred using the DADA2 pipeline^52^, including quality filtering, read merging, chimera removal, and contaminant screening. Taxonomic classification was performed using SILVA v138.2 for 16S rRNA gene sequences^53^, and curated reference databases (BOLD, NCBI nt, and EukRibo) for eukaryotic markers. Non-target sequences (e.g., mitochondria, chloroplasts, non-eukaryotic assignments in COI/18S) were removed prior to downstream analyses.

### Network analysis

Microbial co-occurrence networks were inferred from the 16S dataset using a standardized shared feature space defined as the 200 most abundant ASVs after excluding rare taxa (observed in <2 samples). CLR-transformed relative abundances were used to compute Spearman correlations, retaining edges with |ρ| ≥ 0.7. Network metrics (e.g., components, density, transitivity, modularity, centrality) were calculated using *igraph*. Heatwave-associated changes were assessed using bootstrap subsampling (to control for unequal replication), permutation tests of edge turnover and topology shifts, Fisher Z tests of edge rewiring, and paired Wilcoxon tests of node centrality distributions.

### Further statistical analyses

Statistical analyses were performed in R (v4.5.1) with Rstudio interface (2025.09.1). Visual abundance data were modeled using general linear models (GLMs) with Gamma error distribution and log link, including Treatment, Time (pre-vs post-heatwave), and their interaction. Post hoc contrasts were computed from estimated marginal means with Tukey adjustment and heteroskedasticity-consistent (HC0) robust standard errors. Community composition (visual and metabarcoding datasets) was analyzed using Bray-Curtis dissimilarities and permutational multivariate analysis of variance (PERMANOVA; 999 permutations). Where applicable, Aitchison distance (CLR-transformed counts) was used as a compositional sensitivity analysis. Homogeneity of multivariate dispersion was verified prior to interpretation. Alpha diversity metrics (observed richness, Shannon, Simpson, Chao1) were analyzed using two-way linear models (Time × Treatment) with planned contrasts derived from estimated marginal means. Microbial differential abundance testing was performed using ANCOM-BC2 with Benjamini-Hochberg false discovery rate correction. Metabolic fluxes were analyzed using linear models or t-tests with multiplicity correction, and calcification rates using non-parametric tests (Wilcoxon or Kruskal-Wallis with FDR adjustment). All tests were two-sided. Full code and outputs are available in the online repository.

## Acknowledgements

We are thankful to the numerous members of the Marine Microbiomes Lab for their help during lab- and fieldwork (i.e., BMC preparations, ongoing inoculations), and to the Coastal and Marine Resources Core Lab team for their continuous support, including diving operations, oceanographic equipment, marine permits, and operations. This work was supported by KAUST Competitive Research Grant (CRG) URF/1/4723-01-01.

## Data Availability

Data and code are available upon publication and request.

