## Supplementary Material for "Coral Probiotics Buffer Adjacent Ecosystem-Level Responses to Extreme Marine Heatwave"

### Extended Methods

ARMS Retrieval and Assessment

While ARMS were initially developed as an affordable, non-destructive, and standardized tool for sampling coral reef-associated invertebrates[1], a detailed protocol was later established by Leray and Knowlton (2015)[2] to guide ARMS dismantling and sample processing with the goal of ensuring high-quality DNA preservation of both motile and sessile assemblages. To ensure comparability with classical ARMS studies, we here followed previously established methodological approaches for the morphological and molecular characterization of mobile and sessile organisms[2,3]. However, these procedures were adapted to accommodate the customized biomimetic ARMS deployed in this study, particularly considering size constraints imposed by the experimental treatments and reef patches.

All ARMS were retrieved via SCUBA diving. To prevent the loss of colonizing motile organisms, a transparent zip-lock bag was placed over the ARMS and closed at the metal stick using a cable tie prior to their removal from the benthos. Upon retrieval and once on board, ARMS were maintained submerged in aerated freshly collected seawater from the site during transportation. Zip-lock bags were carefully opened only upon arrival at the research vessel avoiding content mixture or loss of motile invertebrates. Postprocessing (see below) took place at the wet laboratory facilities at KAUST, Saudi Arabia, less than 3 hours after retrieval. Postprocessing started with a non-invasive 1) quantification of biogeochemical functioning of the biomimetic ARMS community ($n = 4$ for each timepoint and treatment), followed by 2) identifying the motile invertebrate associated with each ARMS as well as determining the sizes of collected and identified individuals ($n = 4$ for each timepoint and treatment), 3) disassembling the ARMS structures for taxonomic identification of the sessile invertebrate and photoautotrophic fraction (via photo analysis ($n = 4$ for each timepoint and treatment), and 4) final processing of ARMS for molecular analyses ($n = 4$ for each timepoint and treatment, except for control post heatwave due to technical issues). Details of each procedure are described in detail further below.

*Quantification of Biogeochemical Community Functioning*

To assess the functioning of the ARMS-associated assemblages, incubations to measure fluxes of dissolved oxygen ($O_2$) and total alkalinity (TA) were performed. For this, the metal sticks were cut 1 cm under the lower tile to reduce the size of each structure before placing the entire unit in a PVC jar (2 L volume, Sunpet India, Mumbai, India), including the water that was sampled with each unit. The transparent jar was taped with commercial black tape to ensure no external light influx. Instead of closing the jar with a lid, the incubation flow-sensor head of the 'Community *In-Situ* Metabolism' device (CISME, Qubit Systems, Canada)[4] was used to close the jar airtight. The jar was filled with seawater and was entirely submerged in a water bath that was temperature controlled. Incubation temperatures were set at ambient temperatures measured at the respective sampling time (30.5°C and 26°C, respectively, at pre- and post-marine heatwave sampling points). The flow-sensor head of the CISME consists of an $O_2$ optode for continuous oxygen measurements, a custom pump, a custom LED light source as well as a port that can be manually closed and opened (we refer to Dellisanti et al. (2020)[4] for a detailed overview of technical specification). The incubations started with the port being open for approximately 30 seconds until all air bubbles were outside the system. Incubations for continuous $O_2$ flux measurements were performed for 20 minutes with 45-minute dark-acclimated ARMS under completely dark conditions, followed by 20 minute incubations under daylight conditions (respective ambient temperature at sampling time, light was set to equal photon flux of ~ 400 µM quanta $m^{-2}$ $s^{-1}$ at sampling depth, salinity 39 PTU, pump at '7500') using freshly collected seawater from the reef site. Discrete water samples for TA were taken at the beginning and the end of each incubation. Changes in TA between the start and the end of incubations (both light and dark) were then used to calculate rates of net calcification. Note, changes in seawater chemistry (i.e., dissolved inorganic/organic nitrogen, dissolved organic carbon, phosphate) between start and end of incubations were negligible (see online repository) and therefore not considered when calculating rates. Cryptobenthic community calcification was calculated by concentration differences in TA, which are primarily caused by calcification and dissolution of $CaCO_3$, whereby TA is reduced (increased) by two molar equivalents for every mole of $CaCO_3$ produced (dissolved)[5]. Additionally, blank seawater incubations under the same settings were performed to account for planktonic background metabolism. All rates/fluxes were standardized to the surface area of the ARMS, background metabolism and time taking the incubation jar volume into account. $P_{net}$ and $R_{dark}$ were calculated based on continuous $O_2$ measurements with dissolved $O_2$ sensors from light and dark incubations, respectively. For this, we used the 'auto_rate' function of the respR package[6] in R.studio (version 4.5.1)[7]. $P_{gross}$ was calculated as $P_{gross} = P_{net} + |R_{dark}|$. All rates are expressed as mmol $O_2$ $m^{-2}$ $d^{-1}$ for dissolved oxygen fluxes and mmol $CaCO_3$ $m^{-2}$ $d^{-1}$ for calcification, respectively. All data are reported as mean ± standard error and are reported in the online repository.

*Visual Biodiversity Assessment*

Following the incubations, ARMS were carefully removed from the jars for further analysis. The incubation water was filtered through 2-mm, 500-μm, and 106-μm sieves to retain any mobile specimens. The ARMS were further carefully rinsed with previously filtered seawater from the study area through a 0.2 μm Isopore® membrane before disassembling the tiles. Mobile specimens retained in the sieves were identified alive to the lowest taxonomic level possible based on their morphology using stereo magnifiers (max. 16x magnification) after relaxation of the individuals using a solution of Magnesium Chloride ($MgCl_2$) or, in the case of Brachyura, clove oil. All specimens were measured to determine their size (see online repository for raw data), and subsequently preserved in 95% ethanol. After collecting all unattached individuals, both tiles were photographed from both sides using a Canon R5 camera with external flashes. All images were analyzed by overlaying a 7 × 7 cm digital quadrat subdivided into a 10 × 10 grid, with four fixed points per cell, resulting in a total of 40 stratified points. Presence-absence was assessed at each point to estimate total biotic coverage and the relative proportion of major taxa using PhotoQuad software[8]. After the photographs were taken, the ARMS were placed in trays with filtered seawater and carefully inspected to collect remaining mobile organisms. When still present, mobile organisms were collected for identification and preservation. Sessile organisms and any other biological material attached to the tiles were subsequently removed using a stainless-steel scraper. The resulting material was preserved in 50 ml Falcon tubes with 98% ethanol and stored at +4 °C until further processing (molecular analyses).

*DNA extractions and biological markers (16S rRNA, 18S rRNA and COI genes) sequencing*

Genomic DNA from the scrapings of the miniARMS from both sampling points was extracted using the DNeasy PowerLyzer PowerSoil Kit (Qiagen, Germany). Prior to extraction, samples were air-dried to remove residual ethanol and then homogenized with a mortar and pestle to ensure an even representation of the microbial and eukaryotic communities. Extractions were performed following the manufacturer's protocol with the following modifications depending on the target gene: a) for the 16S rRNA gene, 0.7 g of homogenized material was added to Solution C1 and subjected to a 10-minute shaking at 2000 rpm; b) for the 18S rRNA and cytochrome c oxidase subunit I (COI) genes, 0.7 g of homogenized material was added to Solution C1 with proteinase K and incubated overnight for approximately 14 hours at 56 °C with constant agitation (400 rpm) in a Thermomixer (ThermoFisher®). The remaining steps were carried out following the manufacturer's instructions. DNA samples were stored at - 20°C until downstream analyses. DNA concentration for all samples were quantified using Qubit 4

Fluorometer, with the High Sensitivity DNA Kit (Invitrogen™), and finally, total DNA was normalized to 5 ng/μl.

PCR amplifications and sequencing were performed by Biomarker Technologies (BMKGENE). In summary, PCR amplification for the V3-V4 region of the 16S rRNA gene was carried out using a nested approach. The first round employed primers 243F (5′-GGATGAGCCCGCGGCCTA-3′) and A3R (5′-CCAGCCCCACCTTCGAC-3′) for 10 cycles of 94 °C for 30 s, 56 °C for 40 s, and 72 °C for 90 s, followed by a final extension at 72 °C for 10 min. The second round used barcoded primers S-D-Bact-0341-b-S-17-ad (5′-CCTACGGGNGGCWGCAG-3′) and S-D-Bact-0785-a-A-21-ad (5′-GACTACHVGGGTATCTAATCC-3′), with an initial denaturation at 98 °C for 1 min, followed by 20 cycles of 98 °C for 10 s, 54 °C for 30 s, and 72 °C for 60 s, and a final extension at 72 °C for 5 min. The COI gene was amplified using primers mlCOlintF and HCO2198R under standard conditions, and the V4 region of the 18S rRNA gene was amplified using primers Uni18S and Uni18SR (95 °C 5 min; 30 cycles of 95 °C 30 s, 50 °C 30 s, 72 °C 40 s; final extension 72 °C 5 min). Additionally, null template PCRs (no template input) and negative controls from the DNA extraction were included to account for laboratory contaminants. Amplicons were then purified using VAHTS DNA Clean magnetic beads (0.8× ratio). Library indexing was prepared in 20μL reactions using Q5 High-Fidelity Master Mix and indexing primers with 10 PCR cycles (98 °C 10 s, 65 °C 30 s, 72 °C 30 s), followed by a final extension at 72 °C for 5 min. Indexed libraries were assessed for size distribution and concentration. Libraries were then normalized to equal DNA amounts, pooled in equimolar concentrations, and purified prior to sequencing. Clean libraries were sequenced using NovaSeq6000 platform (Ilumina) employing a paired-end 250 bp strategy.

*Bioinformatic analyses of amplicon sequence libraries*

The standard DADA2 pipeline[9] was used to process the 16S rRNA, 18S rRNA and COI gene-based amplicon libraries. To process the COI amplicon libraries, raw reads were decontaminated of phiX and adapter-trimmed using the BBDuk tool from the BBMap suite (Bushnell B, http://sourceforge.net/projects/bbmap/). PCR primers were removed from the reads using the 'cutadapt' tool[10]. The processed reads were analyzed on DADA2 under the standard inference mode to define the amplicon sequence variants (ASVs). Forward and reverse reads were merged with the *mergePairs* function of DADA2. Chimeric ASVs were identified with the *removeBimeraDenovo* function and removed. All potential contaminant ASVs that were identified by the 'decontam' tool[11] from the DNA extraction blanks or negative control libraries were removed from the analyses. ASVs representing putative pseudogenes or nuclear mitochondrial DNA segments (nuMTs) were identified and filtered out based on length as described in Porter & Hajibabei (2021)[12]. The 'BOLDigger3' tool (Buchner & Leese, 2020)[13] along with 'BLASTn'[14] coupled with the 'IcaPident' function in the biohelper R package (https://github.com/olar785/biohelper/) were used to assign taxonomy to these chimera-, contaminant- and nuMT-free ASV sequences (BLASTn parameters: 5 nt hits per ASV, e-value of 1e-10) based on the curated BOLD database v5[15] as well as the NCBI core nucleotide (nt) database (version 5; January, 2025). Non-target ASVs representing taxa that were outside of Eukaryota were excluded.

For the 16S rRNA and 18S rRNA gene-based amplicon libraries, a similar pipeline as above was followed with the exception of the nuMTs filtering step. 'BLASTn'[14] coupled with the 'IcaPident' function in the 'biohelper' R package (https://github.com/olar785/biohelper/) was used to assign taxonomy to the 18S rRNA ASV sequences (BLASTn parameters: 5 hits per ASV, e-value of 1e-10) based on the EukRibo database[16] and the NCBI core nucleotide (nt) database (version 5; January, 2025). In case of the 16S rRNA ASV sequences, the SILVA database, version 138.2[17] was used at the assignTaxonomy step in the DADA2 workflow. Additionally, the ASV sequences from all samples were queried for the presence of the 16S rRNA gene sequences from each of the bacterial strains in BMC1 and BMC2 using BLASTn (at minimum sequence identity of 97% and query coverage of 90%).

<u>Statistical Analyses and Reproducibility</u>

All statistical analyses were carried out in R (v4.5.1)[7] with the R.studio interface (2025.09.1). We describe the respective analysis below matching the sections presented in the results targeting comparisons across treatments within sampling points (e.g., control vs BMC1 vs BMC2 prior to the MHW), and across sampling points within treatments (e.g., control pre-heatwave vs control post-heatwave). A large language model (ChatGPT) was used to clean the final R codes, and all R codes can be found in the online repository.

*Visual Diversity Analysis*

Biodiversity data of visually identified mobile invertebrates were handled and analyzed using the 'dplyr'[18], 'emmeans'[19], and 'sandwich'[20] packages. Abundances of visually identified motile cryptobenthic macrofauna were standardized to abundance $m^{-2}$ and summed per independent replicate (i.e., ARMS unit) for each treatment × time combination prior to analysis. Total abundance per replicate was analyzed using a generalized linear model (GLM) with a Gamma error distribution and log link to account for strictly positive, right-skewed response data. The model included treatment (Control, BMC1, BMC2), year (pre-, post-heatwave), and their interaction as fixed effects. Post hoc pairwise comparisons among all treatment × year combinations were computed from estimated marginal means (EMMs) on the response scale. To account for potential heteroskedasticity and low replication, heteroskedasticity-consistent (HC0) robust standard errors were applied using a sandwich covariance estimator. *P*-values were adjusted for multiple comparisons using Tukey's method. Reported contrasts (Supplementary Table S1) represent abundance ratios derived from back-transformed model estimates. Temporal changes in total abundance within each treatment (pre- vs post-heatwave) were assessed using treatment-specific generalized linear models with a quasi-Poisson distribution and log link to account for overdispersion in count-link data. From these models, the year coefficient (pre- relative to post-heatwave) was extracted on the log scale along with its standard error and *p*-value. Descriptive mean ± standard error values per treatment and year were calculated from replicate totals, and proportional changes (%) was computed to aid interpretation (Supplementary Table S2).

*Sessile and photoautotrophic fraction*

The sessile and photoautotrophic fraction was taxonomically identified as described above, and all analyses were conducted at the ARMS replicate level, treating each ARMS unit as an independent biological replicate. Data processing and statistical analyses were performed in R using the packages 'tidyverse'[18], 'rstatix'[21], and 'vegan'[22]. For each replicate, species-level abundances were first averaged within each ARMS × Year × Treatment × Species combination. Replicate-level relative abundances were then calculated by normalizing species abundances within each ARMS unit such that row sums equaled one (i.e., species abundance divided by total abundance per replicate). This normalization ensured comparability of community composition across treatments and sampling points independent of total abundance. All subsequent univariate and multivariate analyses were based on these replicate-level relative abundance values.

To test treatment-associated differences within each sampling point (pre- and post-heatwave, respectively), analyses were performed separately for each Species × Year combination. For each species within each year, treatment effects (Control, BMC1, BMC2) were assessed using a Kruskal-Wallis rank-sum test applied to replicate-level relative abundances. To control multiple testing, Benjamini-Hochberg (BH) false discovery rate correction[23] was applied within each year across species, yielding adjusted global p-values. A gatekeeping framework was implemented to reduce false-positive inference: pairwise comparisons were conducted only for Species × Year combinations with raw Kruskal-Wallis *p*-values ≤ 0.10. For these cases, pairwise Wilcoxon rank-sum tests were performed between treatment groups, and Benjamin-Hochberg (BH)-correction was applied within each Species × Year across pairwise contrasts.

Significant pairwise differences were defined at adjusted *p*-values ≤ 0.10. For contextual interpretation, mean relative abundance per treatment was calculated and reported, and significant contrasts were annotated according to the direction of higher relative abundance. A complete unfiltered results table is provided in the online repository, and a compact version restricted to Species × Year combinations with global Kruskal-Wallis $p \leq 0.10$ is presented as Supplementary Table S7.

Community-level differences in sessile assemblage composition were evaluated using multivariate analyses based on Bray-Curtis dissimilarities. For each ARMS × Year × Treatment replicate, a species-by-replicate matrix was constructed using replicate-level relative abundances, with species absent from a replicate assigned zero values. For each year separately, treatment effects on community composition were assessed using permutational multivariate analysis of variance (PERMANOVA) implemented via the 'adonis2' function in the 'vegan' package[22], based on Bray-Curtis dissimilarities with 999 permutations. To evaluate the assumption of homogeneous multivariate dispersion, distances to group centroids were calculated using 'betadisper', followed by permutation testing (999 permutations). $R^2$ values, F statistics, and permutation-based *p*-values are reported in Supplementary Table S8.

*Amplicon metabarcoding community analyses (18S, COI, 16S)*

For metabarcoding datasets (18S rRNA, COI, and 16S rRNA), count tables (ASVs), taxonomy tables, and sample metadata were imported into 'phyloseq'[24] and metadata fields were standardized to match the manuscript factor structure (Time: *Pre Heatwave*, *Post Heatwave*; Treatment: *Control*, *BMC1*, *BMC2*). To reduce noise from extremely rare features, ASVs were filtered prior to downstream analyses by removing singletons/near-singletons (total reads across the full dataset ≤ 1; implemented as 'taxa_sums > 1'). For bacterial 16S analyses, non-target contaminants were excluded by retaining only taxa annotated as Bacteria and removing mitochondria and chloroplast assignments. Following removal of non-target sequences and extremely rare ASVs (total abundance ≤ 1 across the dataset), 4,349 out of initial 4,532 bacterial ASVs were retained for 16S analyses, and 6,090 (out of initial 6,229) and 5,148 (out of initial 5,150) ASVs were retained for the 18S and COI datasets, respectively.

*Beta diversity and PERMANOVAs*

Community-level differences were evaluated using Bray-Curtis dissimilarities computed on relative abundances using the 'vegan' package in R[22]. For 18S and COI, datasets were rarefied to even sequencing depth prior to ordination-free hypothesis testing (18S: 45,000 reads per sample; COI: 47,000 reads per sample; Supplementary Figures S6 and S6), using 'rarefy_even_depth' in the 'phyloseq' package[24], without replacement and a fixed random seed to ensure reproducibility. After rarefaction, counts were converted to relative abundances (expressed as percentages) and Bray-Curtis distances were calculated. Treatment and time effects were tested using PERMANOVA implemented with 'adonis2' in the 'vegan' package (Anderson 2001), with 999 permutations fitting the model distance ~ Time * Treatment. For 18S and COI marker genes, PERMANOVA terms were evaluated sequentially ("by = terms", Type I sums of squares), and results are reported as pseudo-F statistics, partial $R^2$, and permutation *p*-values for Time, Treatment, and their interaction (Time × Treatment), see Supplementary Table S4.

For 16S beta diversity (see Supplementary Table S8), Bray-Curtis analyses followed the same framework, with rarefaction to 60,000 reads per sample prior to calculating relative abundances and Bray-Curtis dissimilarities (Supplementary Fig. S7). In addition, a compositional-distance sensitivity analysis was performed using Aitchison distance (Aitchison 1982), calculated as Euclidean distance on centered log-ratio (CLR) transformed counts. CLR transformation was applied to the cleaned, non-rarefied count table using a pseudo count of 1 to accommodate zeros, and Euclidean distances were then computed on CLR values. Global PERMANOVA tests (Time x Treatment, 999 permutations) were conducted for both Bray-Curtis and Aitchison distance matrices, with targeted one-factor contrasts additionally evaluated for Bray-Curtis where relevant (treatment effects within each time point; and temporal effects within each treatment), again using 'adonis2' with 999 permutations and reporting pseudo-F, $R^2$, and permutation *p*-values.

Across all PERMANOVA models, the assumption of comparable multivariate dispersion among groups was assessed using PERMDISP ('betadisper' followed by 'permutest', 999 permutations, 'vegan' package[22] in R). Dispersion checks were performed for grouping by Time and by Treatment (and within-year treatment tests where applicable), and these diagnostics were reported alongside PERMANOVA outputs to support interpretation of location (centroid) effects versus dispersion-driven differences.

*Alpha diversity (Visual, 18S, COI, 16S)*

Alpha diversity was quantified for visually identified assemblages and for rarefied metabarcoding datasets (18S, COI, 16S). Amplicon datasets were rarefied to even sequencing depth (18S: 45,000; COI: 47,000; 16S: 60,000 reads per sample) following singleton and rare-feature pruning as described above. Rarefaction parameters and random seeds were fixed to ensure exact reproducibility, whereas visual datasets were analyzed without rarefaction. Four diversity indices were calculated using 'estimate_richness' in the 'phyloseq' package[24]: Observed richness, Shannon diversity, Simpson diversity, and Chao1 richness. Chao1 was not calculated for visually identified assemblages because density-based abundance data do not provide the integer singleton/doubleton structure required for non-parametric richness estimation.

For each dataset and diversity metric, treatment and time effects were tested using two-way linear models of the form *Metric ~ Time × Treatment*, fitted with 'lm' in R assuming Gaussian error distributions. Gaussian models were used because diversity metrics were continuous at the patch level and residual diagnostics indicated no strong deviations from homoscedasticity; linear models are robust to moderate departures from normality. Statistical inference was based on Type II ANOVA using the 'car' package[25], and F-statistics and associated $p$-values were extracted for the main effects of Time and Treatment and for the Time × Treatment interaction (Supplementary Table S5). Planned contrasts were conducted using estimated marginal means ('emmeans[19]'). Treatment contrasts (Control vs BMC1 vs BMC2) were performed separately within each sampling time and adjusted using Tukey's method for multiple comparisons. Time contrasts (pre- vs post-heatwave) were conducted separately within each treatment without additional multiplicity correction, as only a single planned comparison was performed per treatment. Significant planned contrasts are reported in Supplementary Table S6, and the full contrast output is available in the online repository.

*Microbial Analyses (16S)*

Differential abundance testing for 16S ASVs was conducted using the 'ANCOM-BC2' package in R (implemented in the ANCOMBC R package[26]). Analyses were performed on the cleaned bacterial dataset (bacteria retained; mitochondria and chloroplast removed; singletons removed via 'taxa_sums > 1'). Hypothesis-driven contrasts were run on biologically motivated subsets: (i) treatment effects within each time point (i.e., pre-heatwave and post-heatwave analyzed separately), and (ii) time effects within each treatment (Control, BMC1, BMC2 analyzed separately). Each ANCOM-BC2 run used 'fix_formula' corresponding to the focal factor (Treatment or Time), 'group' set to the same factor, Benjamini-Hochberg correction for multiple testing[23], and an FDR threshold of 0.05 to define significance. Full ANCOM-BC2 output tables for each contrast (including effect estimates and hypothesis test outputs for all ASVs) were exported for the online repository, while the Supplementary Table S10 reports the total number of ASVs tested and the number of ASVs significant at FDR $\leq 0.05$ for each contrast.

*Microbial co-occurrence network inference and hypothesis-driven topology tests (16S)*

Microbial co-occurrence networks were inferred from the 16S dataset using a controlled, shared feature space to ensure that all between-network contrasts were evaluated comparably. A global ASV universe was first defined as the 200 most abundant ASVs across the full cleaned dataset. For each network-inference subset (Time × Treatment), this global set was further filtered by a within-subset prevalence requirement, retaining only ASVs observed in ≥ 2 samples in that subset, which reduces artefacts driven by sparsely observed features while maintaining a consistent candidate pool across comparisons. Counts were transformed using a centered log-ratio (CLR) transform with a pseudo count of 1 (applied per sample), and pairwise associations were estimated as Spearman correlations among CLR-transformed ASV profiles. Networks were constructed by thresholding correlations at $|\rho| \geq 0.7$, retaining both positive and negative associations (edge sign preserved), and using absolute correlation values as edge weights for topology metrics. To empirically justify the restriction to the 200 most abundant ASVs, a rank-abundance analysis was performed on the prevalence-filtered (observed in ≥ 2 samples) relative abundance table. ASVs were ranked by mean relative abundance across all samples, and cumulative relative abundance was calculated. The top 200 ASVs present in at least 2 samples represented the dominant fraction of total community abundance (>83%, Supplementary Fig. S3), indicating that this threshold captured the majority of biologically relevant signals while excluding the long tail of rare taxa. This approach balances interpretability and robustness in correlation-based network inference.

Network topology metrics were calculated using the 'igraph' package[27] in R, including node and edge counts, number of connected components, edge density, global transitivity, mean shortest-path length (allowing disconnected graphs), diameter, and modularity (community detection via Louvain where possible, with a greedy fallback). Node-level summaries included degree, normalized betweenness, harmonic closeness (robust to disconnected graphs), and eigenvector centrality computed within connected components and scaled for comparability.

Network change across the marine heatwave was assessed with a suite of hypothesis-driven tests: First, to evaluate robustness to unequal replication and sampling variance in the control group ($n = 4$ pre-heatwave, and $n = 3$ post-heatwave), a bootstrap null was generated from the pre-heatwave control subset by repeatedly resampling $k = 3$ samples with replacement (300 iterations), reconstructing a network for each bootstrap replicate using the same pipeline and parameters, and comparing the observed post-heatwave control network metrics to the bootstrap-derived null distributions (one-sided permutation-style *p*-values computed from the proportion of null values at least as extreme as the observation, with a +1 correction). Second, descriptive edge identity turnover was summarized using Jaccard overlap of edge sets between pre- and post-heatwave networks (computed on undirected ASV pairs). Complementarily, a label-shuffling permutation test was used to test whether observed pre/post-heatwave differences in edge overlap and absolute changes in density and modularity were greater than expected under random reassignment of samples to the two groups (199 permutations; networks reconstructed per permutation using the same shared feature space, prevalence filter, CLR transform, and correlation thresholding). Third, edge-level rewiring was evaluated by computing Spearman correlation matrices (on CLR-transformed values) separately for pre- and post-heatwave subsets within each treatment, and testing per-edge correlation differences using Fisher Z statistics with standard errors based on sample sizes; p-values were adjusted across all tested edges using Benjamini-Hochberg, and the number of edges with FDR < 0.05 was reported per treatment. Finally, node-level reorganization was assessed by comparing centrality distributions for ASVs present in both pre- and post-heatwave networks within each treatment using paired Wilcoxon tests (paired by ASV identity), focusing on degree and eigenvector centrality as summary measures of connectivity and influence. Full simulation outputs (bootstrap distributions and permutation nulls), edge-level Fisher-Z tables, and node-level metric tables were exported for the online repository, with an extracted version available presented in Supplementary Table S11.

*Community Functioning*

Community functioning analyses were conducted in R using the packages 'dplyr' and 'readr'[18], 'rstatix'[21], and 'effectsize'[28]. All tests were performed at the level of independent ARMS units. Oxygen flux data included three biologically relevant metabolic rates: net photosynthesis ($P_{net}$), gross photosynthesis ($P_{gross}$), and dark respiration ($R_{dark}$). To assess temporal changes across the marine heatwave within each treatment, two-sample t-tests assuming equal variances were conducted separately for each flux and treatment (Rate ~ Sampling_Point). Because three temporal comparisons were performed per flux (one per treatment), *p*-values were adjusted using Bonferroni correction within each flux. Adjusted *p*-values and corresponding significance codes are reported in Supplementary Table S12A. To quantify the magnitude and direction of temporal change, effect sizes were calculated as Cohen's d for each Treatment × Flux comparison using the 'effectsize' package[28]. Positive values indicate higher pre-heatwave rates relative to post-heatwave rates, and effect sizes were interpreted using conventional thresholds (small ≥ 0.2, medium ≥ 0.5, large ≥ 0.8). To evaluate treatment differences within each sampling point, one-way analyses of variance (ANOVA) were conducted separately for each flux at each time point (Rate ~ Treatment). When ANOVAs were performed, pairwise differences among treatments were assessed using Tukey's Honestly Significant Difference (HSD) test, which inherently adjusts for multiple comparisons. Tukey-adjusted *p*-values are reported in Supplementary Table S12B.

Calcification rates were analyzed using non-parametric tests due to small sample sizes and potential deviations from normality. To assess temporal changes within treatments (pre- vs post-heatwave), Wilcoxon rank-sum tests were conducted separately for each treatment group. Resulting *p*-values were adjusted across treatments using the Benjamini–Hochberg false discovery rate (FDR) procedure[23]. Raw and adjusted *p*-values are reported in Supplementary Table S13A. To evaluate treatment effects within each sampling point, pairwise Wilcoxon rank-sum tests were conducted comparing Control, BMC1, and BMC2. Benjamini-Hochberg correction was applied within each sampling point across pairwise contrasts. Adjusted p-values are reported in Supplementary Table S13B. All statistical tests were two-sided. Exact sample sizes per group are provided in the corresponding tables.

To evaluate whether variation in net community calcification was associated with the abundance of calcifying taxa, we conducted correlation analyses between calcification rates and (i) crustose coralline algae (CCA) abundance and (ii) pooled abundance of calcifying taxa (defined a priori as CCA, Halimeda, Scleractinia, calcifying molluscs, serpulid/spirorbid polychaetes, barnacles, cheilostome bryozoans, and echinoids). Species-level abundances were aggregated per ARMS × Year × Treatment combination to match the scale of calcification measurements. Associations were assessed using Spearman rank correlations to account for small sample sizes and potential non-normality. Correlations were calculated (i) across all patches combined, (ii) separately within treatments, and (iii) for paired temporal changes (Δ2024–2023) to test whether changes in CCA abundance tracked changes in calcification. No significant correlations were detected in any comparison, indicating that variation in net community calcification was not directly explained by variation in CCA cover or pooled calcifier abundance at the replicate scale (see Supplementary Table S14).

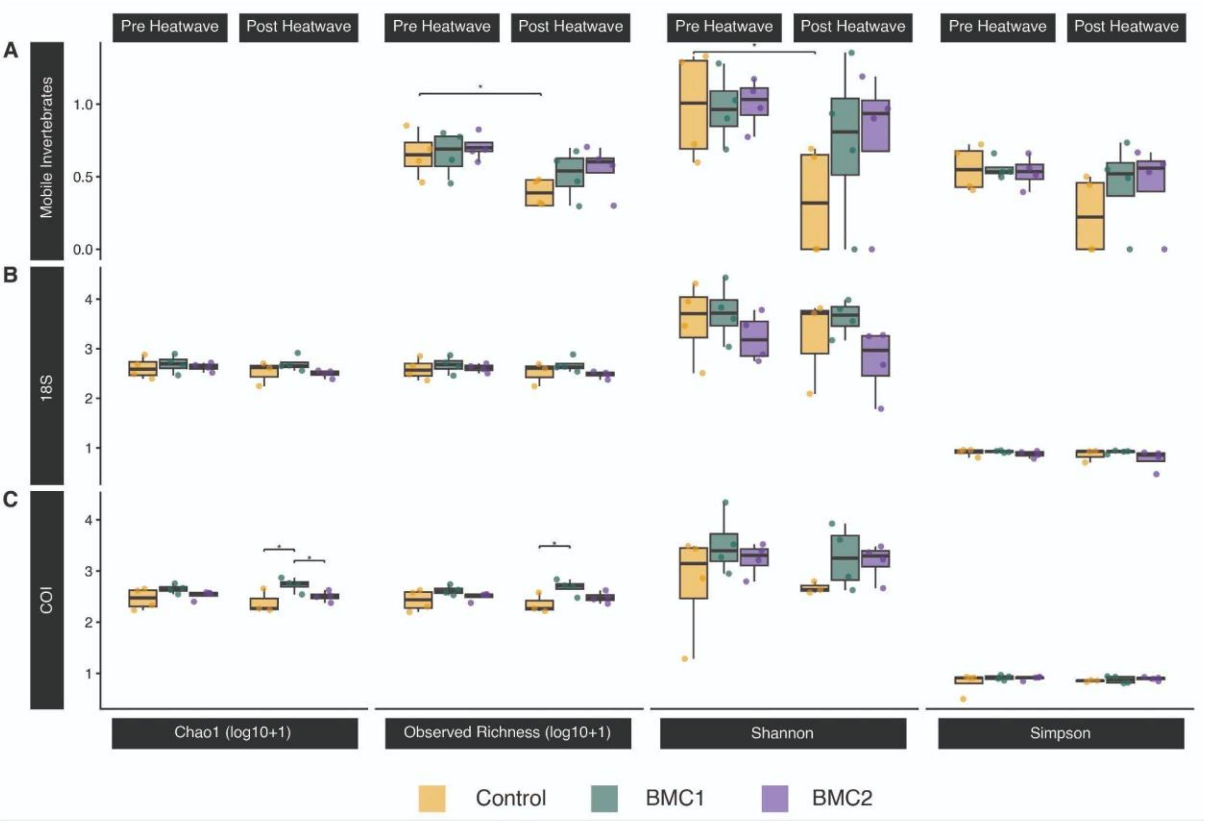

**Supplementary Figure S1: Alpha diversity across visual and metabarcoding datasets before and after the marine heatwave.** Alpha diversity metrics for visually identified mobile invertebrates **(A)**, 18S rRNA gene-derived eukaryotic assemblages **(B)**, and COI-derived metazoan assemblages **(C)** across treatments (Control: gold, BMC1: teal, BMC2: purple) and sampling times (pre- vs post-heatwave). Observed richness and Chao1 richness are shown as log10(value + 1), whereas Shannon and Simpson diversity are displayed on their native scales. Statistical results derive from two-way linear models (Type II ANOVA) testing effects of Time, Treatment, and their interaction (Supplementary Table S5), followed by planned contrasts using estimated marginal means (Supplementary Table S6). Asterisks indicate statistically significant contrasts ($p_{adj} < 0.05$). Visual assemblages showed significant declines in observed richness and Shannon diversity over time in control patches, whereas no time × treatment interactions were detected for metabarcoding datasets. COI-derived metazoan assemblages exhibited significant treatment effects independent of time, with lower post-heatwave richness in control relative to BMC1 patches. No significant interaction effects were detected for any dataset.

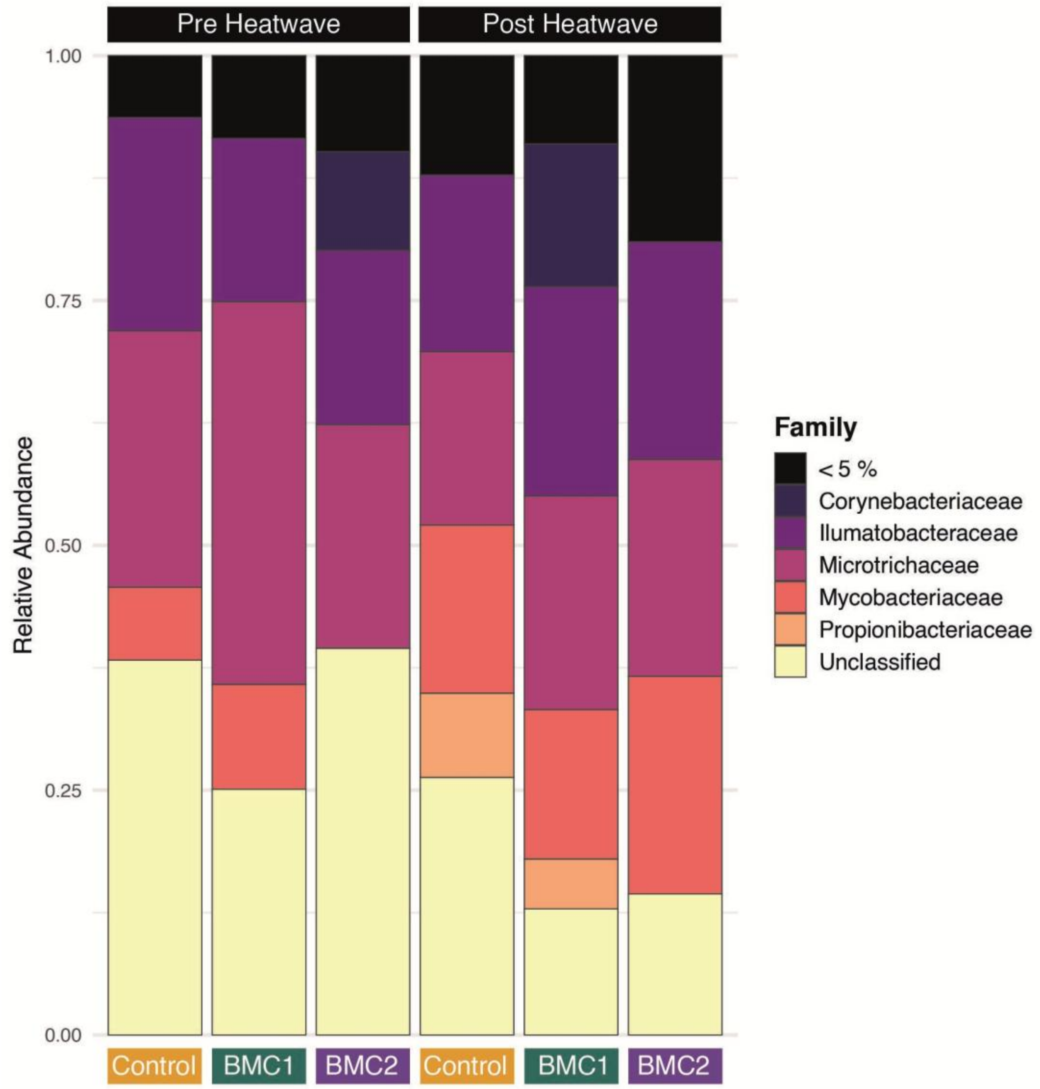

**Supplementary Figure S2: Family-level bacterial community composition across treatments and sampling periods.** Relative abundance of dominant bacterial families associated with cryptobenthic assemblages before and after the 2023 marine heatwave across Control, BMC1, and BMC2 treatments. No consistent treatment- or heatwave-associated shifts in family-level composition were observed, supporting the conclusion that probiotic-associated effects on cryptobenthic microbial communities were not expressed through taxonomic restructuring but rather through changes in interaction network organization.

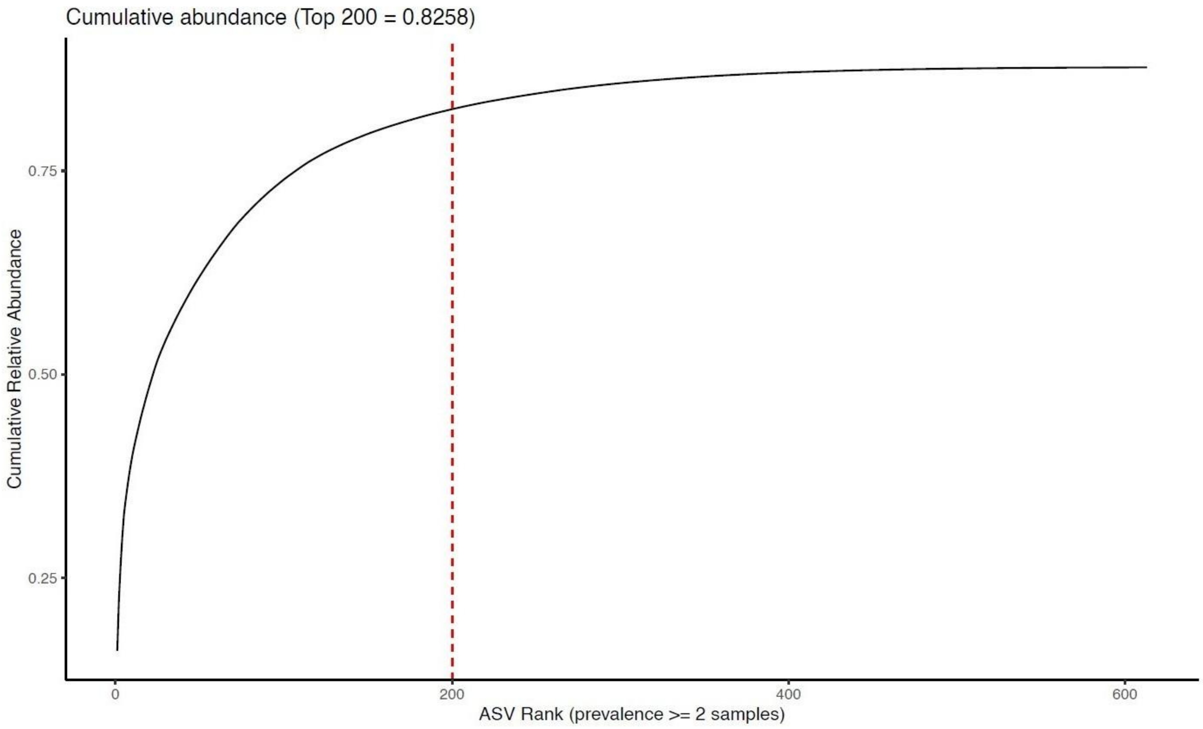

**Supplementary Figure 3: Rank-abundance distribution and cumulative coverage of prevalence-filtered 16S ASVs.** ASVs were first restricted to those observed in ≥ 2 samples across the full dataset to reduce sparsity-driven artefacts. Taxa were then ranked by mean relative abundance (across all samples) and cumulative relative abundance was calculated. The dashed vertical line indicates the threshold used for network inference (top 200 ASVs). The cumulative curve demonstrates that the 200 most abundant ASVs account for the dominant fraction of total community relative abundance (top 200 = 0.826), supporting their use as a standardized, shared feature space for co-occurrence network construction while excluding the long tail of rare taxa.

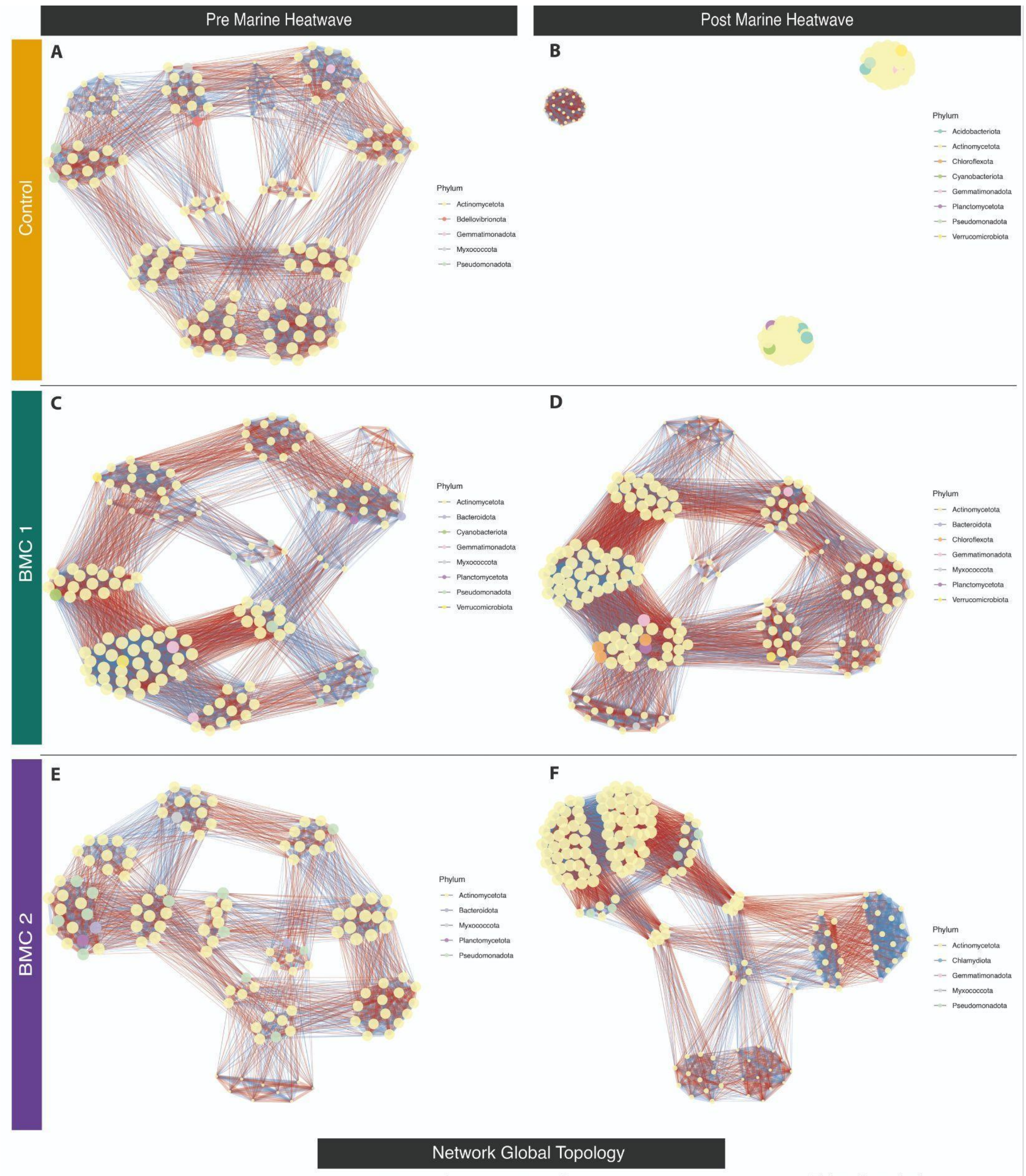

Network Global Topology

| Network | Nodes | Edges | Components | Density | TransGlobal | MeanDist | Diameter | Modularity |
|---|---|---|---|---|---|---|---|---|
| Control (Pre) | 130 | 2824 | 1 | 0.337 | 0.628 | 1.548 | 2.4 | 0.374 |
| Control (Post) | 158 | 4508 | 3 | 0.363 | 1.000 | 1.000 | 1.0 | 0.575 |
| BMC1 (Pre) | 150 | 4273 | 1 | 0.382 | 0.709 | 1.497 | 2.4 | 0.322 |
| BMC1 (Post) | 180 | 5936 | 1 | 0.368 | 0.687 | 1.510 | 2.4 | 0.355 |
| BMC2 (Pre) | 123 | 2498 | 1 | 0.333 | 0.627 | 1.554 | 2.4 | 0.372 |
| BMC2 (Post) | 170 | 5684 | 1 | 0.396 | 0.814 | 1.509 | 2.4 | 0.320 |

**Supplementary Figure S4: Microbial co-occurrence networks constructed from all detected ASVs.** Meta-community co-occurrence networks inferred using all ASVs present in at least two samples are shown for Control **(A-B)**, BMC1 **(C-D)**, and BMC2 **(E-F)** patches before **(A, C, E)** and after **(B, D, F)** the marine heatwave. Networks were constructed using the same inference framework and preprocessing pipeline as in Fig. 4 but without restricting the feature space to the top 200 most abundant ASVs. Broad structural patterns are consistent with those observed in the reduced (Top200 and present in ≥ 2 samples) networks, including pronounced reorganization in control patches following the heatwave and comparatively cohesive architectures in probiotic-treated patches. Inclusion of all ASVs increases network complexity and visual density but does not qualitatively alter the overall interaction trajectories among treatments.

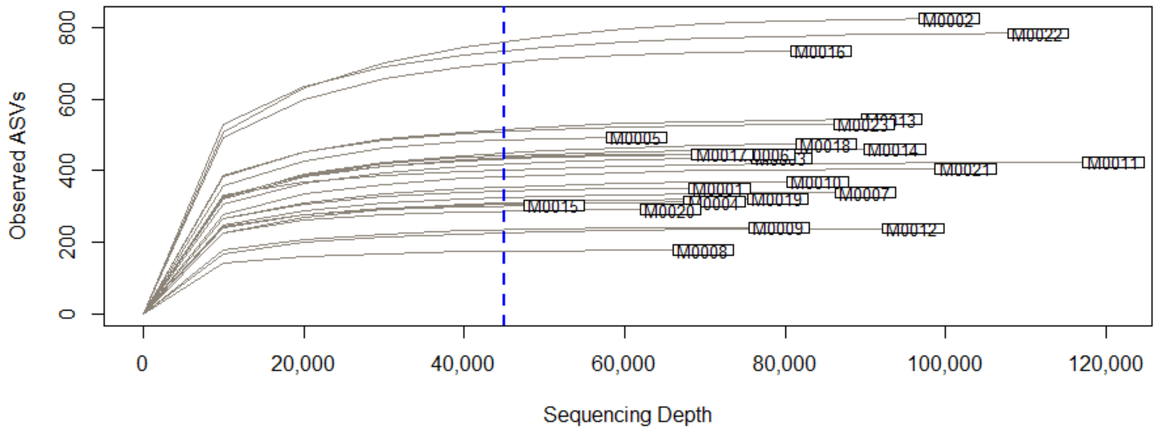

**Supplementary Figure S5: Rarefaction curves (18S rRNA).** Observed ASV richness plotted against sequencing depth for each sample in the cleaned 18S dataset. The dashed blue line marks the rarefaction depth of 45,000 reads applied in downstream diversity analyses. Curves approaching asymptotes indicate sufficient sequencing coverage across samples.

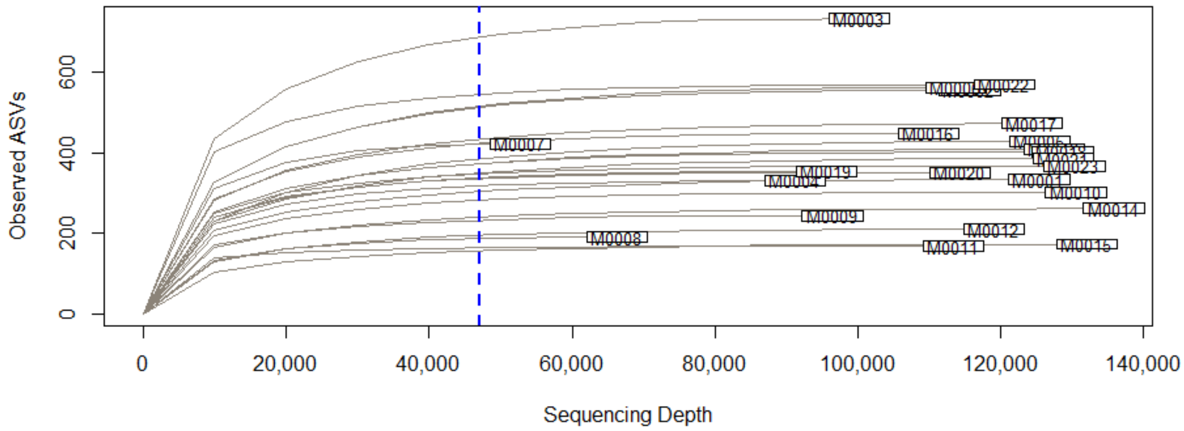

**Supplementary Figure S6: Rarefaction curves (COI rRNA).** Observed ASV richness plotted against sequencing depth for each sample in the cleaned COI dataset. The dashed blue line marks the rarefaction depth of 47,000 reads applied in downstream diversity analyses. Curves approaching asymptotes indicate sufficient sequencing coverage across samples.

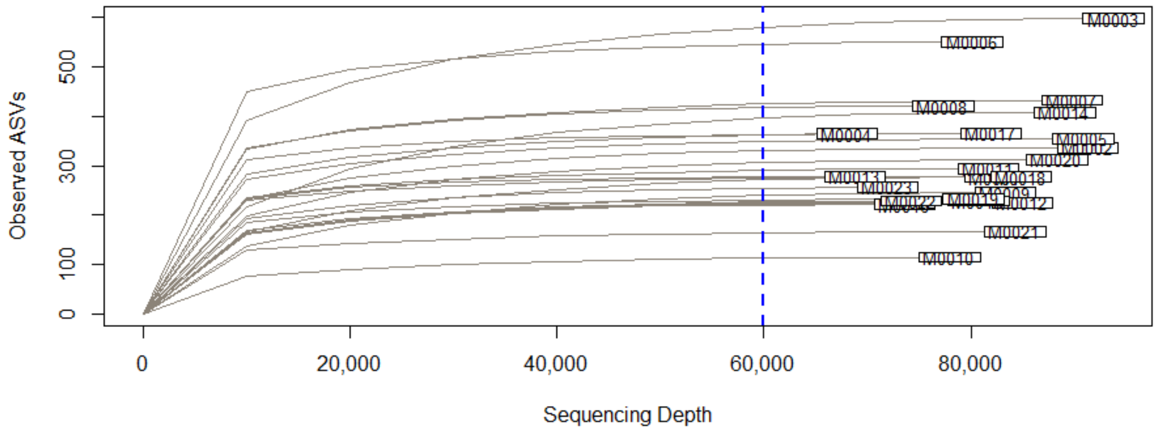

**Supplementary Figure S7: Rarefaction curves (16S rRNA).** Observed ASV richness plotted against sequencing depth for each sample in the cleaned 16S dataset. The dashed blue line marks the rarefaction depth of 60,000 reads applied in downstream diversity analyses. Curves approaching asymptotes indicate sufficient sequencing coverage across samples.

**Supplementary Table S1:** Pairwise post hoc contrasts of visually identified motile cryptobenthic macrofaunal abundance per m² (per ARMS unit) across treatments (Control, BMC1, BMC2) and sampling points (pre-heatwave in 2023; post-heatwave in 2024). Contrasts are based on estimated marginal means from a generalized linear model with gamma error distribution and log link, using heteroskedasticity-consistent (HC0) robust standard errors. Reported values are abundance ratios (treatment × year combinations), with Tukey-adjusted *p*-values. Significant contrasts ($p < 0.05$) indicate higher post-heatwave abundances in probiotic-treated patches relative to controls. Significant *p*-values ($p < 0.05$) are shown in bold.

| **Contrast** | **Ratio** | **SE** | **df** | **t** | ***p*-value** |
|---|---|---|---|---|---|
| BMC1 Pre / BMC2 Pre | 1.000 | 0.340 | 18 | <0.001 | 1.000 |
| BMC1 Pre / Control Pre | 1.092 | 0.496 | 18 | 0.194 | 1.000 |
| BMC1 Pre / BMC1 Post | 1.886 | 0.751 | 18 | 1.595 | 0.612 |
| BMC1 Pre / BMC2 Post | 1.627 | 0.583 | 18 | 1.360 | 0.749 |
| BMC1 Pre / Control Post | 6.385 | 2.409 | 18 | 4.913 | **0.001** |
| BMC2 Pre / Control Pre | 1.092 | 0.443 | 18 | 0.217 | 1.000 |
| BMC2 Pre / BMC1 Post | 1.886 | 0.643 | 18 | 1.862 | 0.455 |
| BMC2 Pre / BMC2 Post | 1.627 | 0.478 | 18 | 1.659 | 0.573 |
| BMC2 Pre / Control Post | 6.385 | 2.022 | 18 | 5.854 | **0.002** |
| Control Pre / BMC1 Post | 1.727 | 0.786 | 18 | 1.200 | 0.831 |
| Control Pre / BMC2 Post | 1.490 | 0.627 | 18 | 0.947 | 0.928 |
| Control Pre / Control Post | 5.846 | 2.558 | 18 | 4.036 | **0.009** |
| BMC1_Post / BMC2 Post | 0.863 | 0.310 | 18 | -0.411 | 0.998 |
| BMC1 Post / Control Post | 3.385 | 1.281 | 18 | 3.222 | **0.046** |
| BMC2Post / Control Post | 3.923 | 1.319 | 18 | 4.064 | **0.008** |

**Supplementary Table S2:** Summary of visually identified phylum-level abundance of motile cryptobenthic macrofaunal communities associated with reef patches before (2023) and after (2024) the 2023 marine heatwave. Mean richness values (± SE) are shown for each treatment (Control, BMC1, BMC2), along with proportional change following the heatwave. Generalized linear model (GLM) coefficients are reported on the log scale with associated standard errors and *p*-values, reflecting treatment-specific changes in richness between years. Mean values are provided for descriptive context; statistical inference is based on GLM estimates and post hoc contrasts reported in Supplementary Table S1.

| Treatment | Mean ± SE (2023) | Mean ± SE (2024) | Change + SE (%) | GLM estimate (log scale) | SE | *p*-value |
|---|---|---|---|---|---|---|
| Control | 113.61 ± 46.89 | 19.43 ± 5.66 | -82.9 ± 8.6 | -1.766 | 0.795 | 0.068 |
| BMC1 | 124.08 ± 40.22 | 65.78 ± 21.42 | -47.0 ± 24.4 | -0.635 | 0.482 | 0.236 |
| BMC2 | 124.08 ± 27.44 | 76.24 ± 19.59 | -38.6 ± 20.8 | -0.487 | 0.343 | 0.205 |

**Supplementary Table S3: Community-weighted mean body size of motile cryptobenthic invertebrates across treatments and years.** Values represent estimated marginal means (EMM ± SE) of family-level community-weighted mean body size, back-transformed to millimeters from linear models fitted to log1p-transformed abundance-weighted size data. Effects were evaluated using a two-way linear model including Treatment, Year, and their interaction. *P*-values correspond to planned within-treatment contrasts comparing post-heatwave (MHW) versus pre-heatwave means, derived from estimated marginal means on the model scale. No significant temporal shifts in community-weighted mean body size were detected within any treatment, indicating that treatment-associated changes in abundance were not accompanied by detectable size-selective restructuring of the community.

| **Treatment** | **EMM ± SE (Pre-MHW)** | **EMM ± SE (Post-MHW)** | ***p*-value** |
|---|---|---|---|
| Control | 4.29 ± 1.47 | 3.31 ± 1.38 | 0.636 |
| BMC1 | 3.80 ± 1.89 | 4.71 ± 1.84 | 0.736 |
| BMC2 | 4.87 ± 1.89 | 2.93 ± 1.09 | 0.362 |

**Supplementary Table S4: Multivariate analyses of eukaryotic community composition based on 18S and COI amplicon sequencing. (A)** Results of sequential permutational multivariate analysis of variance (PERMANOVA; Type I sums of squares) on Bray-Curtis dissimilarities for 18S rRNA and COI marker gene datasets testing the effects of sampling time (pre- vs post-heatwave), treatment, and their interaction on community composition. **(B)** Tests of homogeneity of multivariate dispersion for the same datasets was assessed using PERMDISP (betadisper) with (999) permutations. Significant PERMANOVA effects were not attributable to heterogeneous dispersion, as dispersion did not differ significantly among groups. Significant *p*-values ($p < 0.05$) are shown in bold.

| **(A) Sequential PERMANOVA (Type I sums of squares)** | | | | |
|---|---|---|---|---|
| **Marker Gene** | **Model Term** | **$R^2$** | **F** | ***p*-value** |
| 18S | Time | 0.068 | 1.61 | **0.027** |
| 18S | Treatment | 0.119 | 1.41 | **0.027** |
| 18S | Time x Treatment | 0.093 | 1.10 | 0.224 |
| COI | Time | 0.066 | 1.52 | **0.004** |
| COI | Treatment | 0.101 | 1.15 | 0.079 |
| COI | Time x Treatment | 0.257 | 1.178 | 0.331 |
| **(B) Homogeneity of multivariate dispersion** | | | | |
| **Marker Gene** | **Factor Tested** | **F** | ***p*-value** | |
| 18S | Time | 0.315 | 0.579 | |
| 18S | Treatment | 0.945 | 0.399 | |
| COI | Time | 1.005 | 0.336 | |
| COI | Treatment | 0.093 | 0.906 | |

**Supplementary Table S5: Alpha diversity of visually identified, eukaryotic (18S and COI marker genes), and microbial (16S marker gene) communities associated with reef patches before and after the 2023 marine heatwave.** Results of linear models (Type II ANOVA) testing the effects of sampling time (pre- vs post-heatwave), treatment (Control, BMC1, BMC2), and their interaction on alpha diversity metrics (Observed richness, Shannon diversity, Simpson diversity, and Chao1 richness) derived from visually identified invertebrates and 18S rRNA, COI, and 16S rRNA gene amplicon datasets. For visually identified assemblages, observed richness and Shannon diversity showed significant main effects of Time, indicating reduced diversity following the heatwave, whereas no treatment or time × treatment interaction effects were detected. For eukaryotic metabarcoding datasets (18S and COI), no significant time × treatment interactions were detected for any metric. In COI-derived metazoan assemblages, Treatment significantly influenced observed richness and Chao1 estimates independent of Time. For bacterial (16S) communities, alpha diversity declined modestly across Time (significant main effect of Time for Observed, Shannon, Simpson, and Chao1), but no Treatment or time × treatment interaction effects were detected. Reported values are F-statistics and associated p-values; significant effects ($p < 0.05$) are shown in bold. Chao1 richness was not calculated for visually identified assemblages because density-based abundance data do not provide the integer singleton/doubleton structure required for non-parametric richness estimation.

| **Marker** | **Metric** | **Factor tested** | **F** | ***p*-value** |
|---|---|---|---|---|
| Visual | Observed | Time | 9.677 | **0.006** |
| Visual | Observed | Treatment | 0.895 | 0.426 |
| Visual | Observed | Time x Treatment | 0.315 | 0.734 |
| Visual | Shannon | Time | 5.138 | **0.036** |
| Visual | Shannon | Treatment | 0.745 | 0.489 |
| Visual | Shannon | Time x Treatment | 0.715 | 0.502 |
| Visual | Simpson | Time | 3.495 | 0.078 |
| Visual | Simpson | Treatment | 0.501 | 0.614 |
| Visual | Simpson | Time x Treatment | 0.648 | 0.535 |
| 18S | Observed | Time | 0.832 | 0.374 |

| | | | | |
|---|---|---|---|---|
| 18S | Observed | Treatment | 1.594 | 0.232 |
| 18S | Observed | Time x Treatment | 0.225 | 0.801 |
| 18S | Shannon | Time | 1.260 | 0.277 |
| 18S | Shannon | Treatment | 2.258 | 0.135 |
| 18S | Shannon | Time x Treatment | 0.169 | 0.846 |
| 18S | Simpson | Time | 1.488 | 0.239 |
| 18S | Simpson | Treatment | 1.744 | 0.205 |
| 18S | Simpson | Time x Treatment | 0.376 | 0.692 |
| 18S | Chao1 | Time | 0.855 | 0.368 |
| 18S | Chao1 | Treatment | 1.578 | 0.235 |
| 18S | Chao1 | Time x Treatment | 0.257 | 0.776 |
| COI | Observed | Time | 0.063 | 0.805 |
| COI | Observed | Treatment | 6.495 | **0.008** |
| COI | Observed | Time x Treatment | 0.645 | 0.537 |
| COI | Shannon | Time | 0.295 | 0.594 |
| COI | Shannon | Treatment | 2.513 | 0.111 |
| COI | Shannon | Time x Treatment | 0.066 | 0.936 |
| COI | Simpson | Time | 0.014 | 0.906 |
| COI | Simpson | Treatment | 1.171 | 0.334 |
| COI | Simpson | Time x Treatment | 0.326 | 0.726 |
| COI | Chao1 | Time | 0.114 | 0.740 |
| COI | Chao1 | Treatment | 6.721 | **0.007** |
| COI | Chao1 | Time x Treatment | 0.779 | 0.474 |
| 16S | Observed | Time | 4.673 | **0.045** |
| 16S | Observed | Treatment | 1.674 | 0.217 |

| | | | | |
|---|---|---|---|---|
| 16S | Observed | Time x Treatment | 0.708 | 0.507 |
| 16S | Shannon | Time | 5.434 | **0.032** |
| 16S | Shannon | Treatment | 1.333 | 0.290 |
| 16S | Shannon | Time x Treatment | 0.053 | 0.949 |
| 16S | Simpson | Time | 7.769 | **0.013** |
| 16S | Simpson | Treatment | 0.709 | 0.506 |
| 16S | Simpson | Time x Treatment | 0.477 | 0.629 |
| 16S | Chao1 | Time | 4.528 | **0.048** |
| 16S | Chao1 | Treatment | 1.635 | 0.220 |
| 16S | Chao1 | Time x Treatment | 0.716 | 0.503 |

**Supplementary Table S6.** Significant planned pairwise contrasts in alpha diversity of visually identified and metabarcoding-derived (16S, 18S, COI) communities associated with reef patches before and after the marine heatwave. Two-way linear models assuming Gaussian error distributions, including sampling time (pre- vs post-heatwave), treatment (Control, BMC1, BMC2), and their interaction, were followed by *a priori* contrasts using estimated marginal means. Treatment contrasts (Control vs BMC1 vs BMC2) were conducted separately within each sampling time and adjusted using Tukey's method for multiple comparisons. Time contrasts (pre vs post) were conducted separately within each treatment without additional multiplicity correction, as only a single planned comparison was performed per treatment. The table reports contrasts with adjusted *p*-values < 0.05. Estimates represent differences between group means (first level minus second level), and confidence intervals correspond to 95% intervals derived from the linear models. Results from all pairwise contrasts in alpha diversity are available in the online repository.

| **Source** | **Metric** | **Contrast** | **Estimate** | **CI low** | **CI high** | $p_{adj}$ |
|---|---|---|---|---|---|---|
| COI | Chao1 | Control Post vs BMC 1 Post | -283.928 | -517.258 | -50.599 | 0.016 |
| COI | Chao1 | BMC1 Post vs BMC2 Post | 229.724 | 13.702 | 445.756 | 0.036 |
| COI | Observed | Control Post vs BMC1 Post | -261.583 | -476.217 | -44.950 | 0.017 |
| Visual | Observed | Control Pre vs Control Post | 2.250 | 0.300 | 5.200 | 0.026 |
| Visual | Shannon | Control Pre vs Control Post | 0.653 | 0.053 | 1.254 | 0.035 |

**Supplementary Table S7: Treatment-associated differences in sessile and photoautotrophic cryptobenthic fractions within each year.** For each sessile category (Taxon) within each sampling year, differences among treatments (Control, BMC1, BMC2) were tested using Kruskal-Wallis (KW) tests on replicate-level relative abundances (H statistic, degrees of freedom, and raw *p*-values shown). Benjamini-Hochberg (BH)-adjusted *p*-values are reported for completeness. For brevity, only Species × Year combinations with raw Kruskal-Wallis $p \leq 0.1$ are shown; complete unfiltered results are available in the online repository. Where this global test criterion was met, pairwise Wilcoxon comparisons were conducted and BH-adjusted p-values are reported; the "higher" label indicates the treatment with the higher mean relative abundance. Complete unfiltered tables for all tested Species × Year combinations, including non-significant results, are provided in an online repository.

| **Sampling Point** | **Taxon** | **KW_H** | ***p*-value** | $p_{adj}$ | **Mean Control** | **Mean BMC1** | **Mean BMC2** | **Sig_pairwise** |
|---|---|---|---|---|---|---|---|---|
| 2023 | Gastropoda | 5.000 | 0.082 | 0.931 | 0 | 0 | 0 | None |
| 2024 | Biofilm | 7.136 | 0.028 | 0.501 | 0.12 | 0.02 | 0.04 | BMC1 < Control (BH p=0.078); BMC2 < Control (BH p=0.078) |
| 2024 | CCA | 6.182 | 0.046 | 0.501 | 0.08 | 0.20 | 0.21 | BMC1 > Control (BH p=0.078); BMC2 > Control (BH p=0.078) |
| 2024 | Serpulinae | 4.599 | 0.1 | 0.623 | 0.04 | 0.03 | 0.01 | None |

**Supplementary Table S8: Multivariate analysis of cryptobenthic assemblages.** PERMANOVA (Bray-Curtis dissimilarities; 999 permutations) and tests of homogeneity of multivariate dispersion (PERMDISP; 999 permutations) assessing treatment effects (Control, BMC1, BMC2) on sessile and photoautotrophic cryptobenthic assemblages within each year. PERMANOVA results report $R^2$, F-statistics, and permutation-based *p*-values; PERMDISP tests evaluate differences in within-group dispersion.

| **Year** | **Method** | **Factor** | **$R^2$** | **F** | ***p*-value** |
|---|---|---|---|---|---|
| 2023 | PERMANOVA (Bray-Curtis) | Treatment | 0.049 | 0.208 | 0.997 |
| 2023 | PERMDISP | Treatment | - | 0.445 | 0.654 |
| 2024 | PERMANOVA (Bray-Curtis) | Treatment | 0.189 | 0.932 | 0.516 |
| 2024 | PERMDISP | Treatment | - | 0.689 | 0.590 |

**Supplementary Table S9: Beta diversity analyses of bacterial communities associated with cryptobenthic assemblages. (A)** Results of permutational multivariate analyses of variance (PERMANOVA) based on Bray-Curtis dissimilarities and Aitchison (CLR-Euclidean) distances of 16S rRNA gene amplicon data. Summary of global models testing the effects of sampling time (pre- vs post-heatwave), treatment (Control, BMC1, BMC2), and their interaction. **(B)** Targeted PERMANOVA contrasts based on Bray-Curtis dissimilarities within time points and within treatments across time, conducted to interpret the significant global Bray-Curtis interaction detected in panel A. Aitchison-based post hoc contrasts are not shown because the corresponding global interaction was not significant. Bray-Curtis analyses detected a significant but diffuse time × treatment interaction, while Aitchison-based analyses did not detect significant interaction effects. Homogeneity of multivariate dispersions was met for all tests. Significant *p*-values are shown in bold ($p < 0.05$).

| **(A) Global PERMANOVA (Time x Treatment)** | | | | | |
|---|---|---|---|---|---|
| **Distance metric** | **Source of Variation** | **DF** | **$R^2$** | **F** | ***p*-value** |
| Bray-Curtis | Model | 5 | 0.325 | 1.639 | **0.004** |
| Bray-Curtis | Residual | 17 | 0.675 | NA | NA |
| Bray-Curtis | Total | 22 | 1 | NA | NA |
| Aitchinson | Model | 5 | 0.233 | 1.031 | 0.298 |
| Aitchinson | Residual | 17 | 0.767 | NA | NA |
| Aitchinson | Total | 22 | 1 | NA | NA |
| **(B1) Treatment effects within time points** | | | | | |
| **Contrast** | **DF** | **$R^2$** | **F** | ***p*-value** | |
| Pre MHW (2023) | 2 | 0.190 | 1.053 | 0.356 | |
| Post MHW (2024) | 2 | 0.171 | 0.827 | 0.919 | |
| **(B2) Temporal effects within treatments** | | | | | |
| **Contrast** | **DF** | **$R^2$** | **F** | ***p*-value** | |
| Control | 1 | 0.307 | 2.232 | 0.071 | |
| BMC1 | 1 | 0.171 | 1.236 | 0.171 | |
| BMC2 | 1 | 0.319 | 2.811 | **0.025** | |

**Supplementary Table S10: Differential abundance testing of bacterial ASVs using ANCOM-BC2.** Differential abundance analysis of 16S rRNA gene amplicon data from bacterial communities associated with cryptobenthic assemblages. ANCOM-BC2 was used to test for differences among treatments (Control, BMC1, BMC2) within each sampling period **(A)** and for temporal shifts (pre- vs post-heatwave) within each treatment **(B)**. P-values were adjusted using the Benjamini-Hochberg false discovery rate (FDR). No ASVs were identified as significantly differentially abundant after correction (FDR ≤ 0.05) in any contrast. Full ANCOM-BC2 output tables are provided in the online repository.

| **(A) Treatment effects within time points** | | | |
|---|---|---|---|
| **Time Point** | **Contrast** | **ASVs tested** | **Significant ASVs (FDR ≤ 0.05)** |
| Pre-MHW (2023) | Treatment | 333 | 0 |
| Post-MHW (2024) | Treatment | 414 | 0 |
| **(B) Time effects within treatments** | | | |
| **Treatment** | **Contrast** | **ASVs tested** | **Significant ASVs (FDR ≤ 0.05)** |
| Control | Pre vs Post | 1361 | 0 |
| BMC1 | Pre vs Post | 2174 | 0 |
| BMC2 | Pre vs Post | 1651 | 0 |

**Supplementary Table S11: Network robustness and topology-change tests of cryptobenthic microbial co-occurrence networks.** Summary of hypothesis-driven tests assessing robustness, similarity, and reorganization of microbial co-occurrence networks across the marine heatwave. Network analyses were inferred using a globally defined set of the 200 most abundant ASVs across the full dataset, further restricted to ASVs observed in ≥ 2 samples within each network inference subset. This ensured that all contrasts were evaluated within a shared feature space and minimized artefacts driven by sparsely observed taxa. Reported tests include: i) bootstrap robustness analysis comparing the observed post-heatwave control network against a null distribution generated by reconstructing networks from bootstrap-resampled subsets of pre-heatwave control samples (k = 3, with replacement); ii) edge- density contrasts (Jaccard overlap) and label-shuffling permutation tests for differences in density/modularity; and iii) node-level permutation tests comparing centrality distributions across pre/post states. Edge-level Fisher-Z tests detected no individual ASV-ASV associations with significant changes in correlation strength in Controls after FDR correction, indicating that topological shifts reflect distributed system-level restructuring rather than isolated rewiring events. Full simulation outputs, permutation distributions, and edge-level statistics are available in the online repository.

| **Treatment** | **Test Category** | **Network metric** | **Test description** | **Statistic/Effect** | ***p*-value** | **Interpretation** |
|---|---|---|---|---|---|---|
| Control | Bootstrap robustness | Nodes | Post vs bootstrap null (Pre-Control, k=3 with replacement) | Observed above null (100) | 0.003 (upper-tail) | Post-Control retains more prevalent taxa than expected under Pre-only sampling |
| Control | Bootstrap robustness | Edge count | Post vs bootstrap null | Observed **above** null (1683) | 0.003 (upper-tail) | Post-Control has more retained associations than expected under Pre-only sampling |
| Control | Bootstrap robustness | Edge density | Post vs bootstrap null | Observed **below** null (0.34) | 0.003 (lower-tail) | Despite more nodes/edges, Post-Control is globally less dense than expected |

| | | | | | | |
|---|---|---|---|---|---|---|
| Control | Bootstrap robustness | Modularity | Post vs bootstrap null | Observed **above** null (0.623) | 0.003 (upper-tail) | Strong increase in compartmentalization relative to Pre bootstrap expectation |
| Control | Network similarity (descriptive) | Jaccard overlap | Pre vs Post edge overlap | J = 0.098 | - | Moderate overlap; edge identities differ despite shared node universe |
| Control | Permutation contrast (label shuffle) | Jaccard overlap | “Low overlap vs null?” | J = 0.098 | 0.005 | Observed overlap not lower than permuted null under this test framework |
| Control | Permutation contrast (label shuffle) | Modularity difference | - | Δmodularity = 0.208 (0.414→0.623) | 0.005 | Modularity difference significant under label-shuffle framework (small n); bootstrap test is the robust inference |
| Control | Node-level reorganization | Eigenvector centrality | Paired ASV-wise shift | Distribution shift | <0.001 | Central taxa restructured across heatwave |
| BMC1 | Permutation contrast (label shuffle) | Density / Modularity | Pre vs Post | Δdensity= -0.011; Δmodularity = 0.057 | all 0.505 | No detectable global-metric shift in topology |
| BMC1 | Node-level reorganization | Eigenvector centrality | Paired ASV-wise shift | Distribution shift | <0.001 | Centrality structure changes despite stable global metrics |
| BMC1 | Edge-level rewiring | Fisher-Z | Pre vs Post | 4555 edges FDR<0.05 | - | Widespread correlation-strength changes; global metrics remain stable |

| | | | | | | |
|---|---|---|---|---|---|---|
| BMC2 | Permutation contrast (label shuffle) | Edge density | Pre vs Post | Δdensity = +0.096 (0.339→0.435) | 0.335 | Post network becomes denser |
| BMC2 | Permutation contrast (label shuffle) | Modularity | Pre vs Post | Δmodularity = −0.077 (0.368→0.291) | 0.335 | Post network becomes less modular (more integrated) |
| BMC2 | Node-level reorganization | Degree centrality | Paired ASV-wise shift | Distribution shift | <0.001 | Strong redistribution of node connectivity |
| BMC2 | Edge-level rewiring | Fisher-Z | Pre vs Post | 5785 edges FDR<0.05 | - | Extensive rewiring accompanies topology shift |

**Note:** Edge-level Fisher Z tests detected no individual ASV-ASV associations with significant changes after false discovery rate correction, indicating that network reorganization reflects distributed system-wide restructuring rather than isolated rewiring events.

**Supplementary Table S12: Metabolic responses of cryptobenthic communities across the marine heatwave.** Results of metabolic incubations comparing cryptobenthic community fluxes pre and post the marine heatwave (MHW). Net photosynthesis ($P_{net}$), gross photosynthesis ($P_{gross}$), and dark respiration ($Rd_{ark}$) were measured at *in-situ* temperatures for each sampling period. **(A)** Within-treatment comparisons (Pre vs Post) were assessed using independent-samples *t*-tests. *P*-values were adjusted using Bonferroni correction across treatments within each flux. Effect sizes are reported as Cohen's *d* (signed), indicating the direction and magnitude of temporal change. **(B)** Pairwise comparisons among treatments within each sampling period were conducted using one-way ANOVA followed by Tukey's HSD post hoc tests. Reported p-values correspond to Tukey-adjusted comparisons. Significant adjusted *p*-values ($p < 0.05$) are shown in bold. Effect sizes are provided to aid interpretation of biological relevance.

| **(A) Metabolism Comparison with Time (Pre vs Post)** | | | | |
|---|---|---|---|---|
| **Flux** | **Treatment** | ***P*-value (raw)** | **$P_{adj}$** | **Effect size (Cohens *d*)** |
| Pnet | Control | 0.314 | 0.942 | 0.855 |
| Pnet | BMC1 | 0.001 | **0.004** | 4.083 |
| Pnet | BMC2 | 0.272 | 0.816 | 0.856 |
| Rdark | Control | 0.377 | 1 | -0.740 |
| Rdark | BMC1 | 0.107 | 0.321 | -1.338 |
| Rdark | BMC2 | 0.969 | 1 | -0.029 |
| Pgross | Control | 0.288 | 0.865 | 0.908 |
| Pgross | BMC1 | 0.008 | **0.025** | 0.2729 |
| Pgross | BMC2 | 0.545 | 1 | 0.454 |
| **(B) Metabolism comparison among treatments** | | | | |
| **Flux** | **Time** | **Treatment** | **$P_{adj}$** | |
| Pnet | Pre MHW | Control vs BMC1 | 0.665 | |
| Pnet | Pre MHW | Control vs BMC2 | 0.912 | |
| Pnet | Pre MHW | BMC1 vs BMC2 | 0.869 | |
| Pnet | Post MHW | Control vs BMC1 | 0.949 | |

| | | | |
|---|---|---|---|
| Pnet | Post MHW | Control vs BMC2 | 0.849 |
| Pnet | Post MHW | BMC1 vs BMC2 | 0.969 |
| Rdark | Pre MHW | Control vs BMC1 | 1.000 |
| Rdark | Pre MHW | Control vs BMC2 | 0.752 |
| Rdark | Pre MHW | BMC1 vs BMC2 | 0.716 |
| Rdark | Post MHW | Control vs BMC1 | 0.710 |
| Rdark | Post MHW | Control vs BMC2 | 0.929 |
| Rdark | Post MHW | BMC1 vs BMC2 | 0.902 |
| Pgross | Pre MHW | Control vs BMC1 | 0.879 |
| Pgross | Pre MHW | Control vs BMC2 | 0.991 |
| Pgross | Pre MHW | BMC1 vs BMC2 | 0.790 |
| Pgross | Post MHW | Control vs BMC1 | 0.949 |
| Pgross | Post MHW | Control vs BMC2 | 0.996 |
| Pgross | Post MHW | BMC1 vs BMC2 | 0.918 |

**Supplementary Table S13. Treatment-dependent patterns of cryptobenthic calcification before and after the heatwave.** Results of statistical tests assessing differences in net calcification rates of cryptobenthic communities across treatments and sampling periods (pre- and post-marine heatwave (MHW)). Panel **(A)** shows within-treatment comparisons across time, testing for heatwave-associated changes in calcification for each treatment. Panel **(B)** presents pairwise comparisons among treatments within each sampling period. Raw and Benjamini-Hochberg-adjusted *p*-values ($p_{adj}$) are reported. Significant adjusted *p*-values ($p < 0.05$) are shown in bold. Together, these analyses quantify treatment-specific differences in carbonate production and their persistence across the marine heatwave.

| **(A) Calcification Comparison with Time (Pre vs Post)** | | | |
|---|---|---|---|
| **Treatment** | ***P*-value (raw)** | $\mathbf{P_{adj}}$ | |
| Control | 0.029 | **0.043** | |
| BMC1 | 0.029 | **0.043** | |
| BMC2 | 0.343 | 0.343 | |
| **(B) Calcification comparison among treatments** | | | |
| **Time** | **Treatment** | ***p*-value (raw)** | $\mathbf{P_{adj}}$ |
| Pre MHW | Control vs BMC1 | 0.029 | **0.029** |
| Pre MHW | Control vs BMC2 | 0.029 | **0.029** |
| Pre MHW | BMC1 vs BMC2 | 0.029 | **0.029** |
| Post MHW | Control vs BMC1 | 0.029 | **0.043** |
| Post MHW | Control vs BMC2 | 0.029 | **0.043** |
| Post MHW | BMC1 vs BMC2 | 0.886 | 0.886 |

**Supplementary Table S14: Spearman rank correlations between net calcification and sessile calcifier metrics, including CCA cover and pooled calcifier abundance.** Correlations were calculated (i) across all patches (n = 22), (ii) for within-patch changes between 2023 and 2024 (Δ values; n = 11 paired patches), and (iii) separately within each treatment (Control, BMC1, BMC2). In all cases, correlations were weak and not statistically significant (all $p > 0.3$). These results indicate that spatial variation and temporal shifts in net carbonate flux were not explained by CCA cover or overall calcifier (defined a priori as CCA, Halimeda, Scleractinia, calcifying molluscs, serpulid/spirorbid polychaetes, barnacles, cheilostome bryozoans, and echinoids) abundance alone, including within individual treatments, suggesting that treatment-related differences in calcification reflect mechanisms beyond simple changes in calcifier cover.

| **Comparison** | **n** | **Spearman's rho** | ***p*-value** |
|---|---|---|---|
| **All patches** | | | |
| Overall: calcification vs CCA cover | 22 | 0.184 | 0.414 |
| Overall: calcification vs pooled calcifiers | 22 | 0.093 | 0.679 |
| **All patches (paired)** | | | |
| Change (2024-2023): ΔCalcification vs ΔCCA | 11 | -0.082 | 0.818 |
| **Control patches** | | | |
| Calcification vs CCA cover (within treatment) | 6 | -0.486 | 0.356 |
| Calcification vs pooled calcifiers (within treatment) | 6 | -0.429 | 0.419 |
| **BMC1** | | | |
| Calcification vs CCA cover (within treatment) | 8 | 0.357 | 0.389 |
| Calcification vs pooled calcifiers (within treatment) | 8 | 0.429 | 0.299 |
| **BMC2** | | | |
| Calcification vs CCA cover (within treatment) | 8 | 0.000 | 1.000 |
| Calcification vs pooled calcifiers (within treatment) | 8 | 0.095 | 0.840 |
